# Beyond BRCA: Discovery of novel causes and consequences of homologous recombination deficiencies

**DOI:** 10.1101/2022.10.01.510467

**Authors:** Daniel J. McGrail, Yang Li, Roger S. Smith, Bin Feng, Hui Dai, Yongsheng Li, Limei Hu, Briana Dennehey, Sharad Awasthi, Marc L. Mendillo, Gordon B. Mills, Shiaw-Yih Lin, S. Stephen Yi, Nidhi Sahni

## Abstract

Since the discovery of *BRCA1* and *BRCA2* mutations as cancer risk factors, we have gained substantial insight into their role in maintaining genomic stability through homologous recombination (HR) DNA repair. However, upon pan-cancer analysis of tumors from The Cancer Genome Atlas (TCGA), we found that mutations in *BRCA1/2* and other classical HR genes only identified 10-20% of tumors that display genomic evidence of HR deficiency (HRD), suggesting that the cause of the vast majority of HR defects in tumors is unknown. As HRD both predisposes individuals to cancer development and leads to therapeutic vulnerabilities, it is critical to define the spectrum of genetic events that drive HRD. Here, we employed a network-based approach leveraging the abundance of molecular characterization data from TCGA to identify novel drivers of HRD. We discovered that over half of putative genes driving HRD originated outside of canonical DNA damage response genes, with a particular enrichment for RNA binding protein (RBP)-encoding genes. These novel drivers of HRD were cross-validated using an independent ICGC cohort, and were enriched in GWAS loci associated with cancer risk. Experimental approaches validated over 90% of our predictions in a panel of 50 genes tested by siRNA and 31 additional engineered mutations identified from TCGA patient tumors. Moreover, genetic suppression of identified RBPs or pharmacological inhibition of RBPs induced PARP inhibition. Further mechanistic studies indicate that some RBPs are recruited to sites of DNA damage to facilitate repair, whereas others control the expression of canonical HR genes. Overall, this study greatly expands the repertoire of known drivers of HRD and their contributions to DNA damage repair, which has implications for not only future mechanistic studies, but also for genetic screening and therapy stratification.

**HIGHLIGHTS:**

- The majority of HR deficiencies detected cannot be directly attributed to aberrations in canonical HR genes.
- Integrated network analysis identifies RNA binding proteins (RBPs) as a novel driver of HR deficiency in patient tumors.
- RBP dysfunction can produce HR deficiencies through both dysregulation of canonical HR genes and action at sites of DNA damage.

## INTRODUCTION

Genomic instability is a hallmark of cancer (Hanahan and Weinberg, 2011), with implications both for treatment strategies as well as cancer screening and prevention. As normal healthy cells have largely intact DNA damage response (DDR) pathways, therapeutic avenues that selectively target tumor cells with DNA repair defects have emerged as a promising treatment strategy (Pilié et al., 2018). For instance, Poly (ADP-ribose) polymerase (PARP) inhibitors have emerged as a powerful approach to treat patients with defects in *BRCA1* or *BRCA2*, genes involved in homologous recombination (HR) repair, a process that faithfully repairs DNA double strand breaks (DSBs) (Brown et al., 2016; Livraghi and Garber, 2015; Pilié et al., 2018). Similarly, microsatellite instability, caused by defects in DNA mismatch repair (MMR), was recently approved as a biomarker for response to immune checkpoint blockade, marking the first approval of tumor type agnostic biomarker by the U.S. Food and Drug Administration (FDA) (Lemery et al., 2017).

Although these DNA repair defects in either the HR or MMR pathways may provide treatment options, non-functional variants of *BRCA1*/*BRCA2* or genes involved in the MMR pathway are also primary drivers in familial cancers (Garber and Offit, 2005). A large fraction (50%) of known drivers of hereditary cancers are genes involved in DNA repair and genome maintenance (Garber and Offit, 2005). In the case of breast cancer, women with a mother, sister, or daughter with breast cancer have a two-fold higher risk of developing breast cancer (Beral et al., 2001); however, only 15-25% of patients with hereditary breast/ovarian cancers have *BRCA1* or *BRCA2* mutations and the majority of hereditary drivers have not yet been identified (Couch et al., 2014; Nielsen et al., 2016). Incomplete knowledge of genetic risk factors hinders approaches for effective cancer screening and prevention. A better understanding of these genetic risks could identify which patients require more aggressive risk reduction approaches, and minimize use of aggressive risk reduction interventions in patients who lack pre-disposition genes (Nelson et al., 2014).

Thus, there exists a critical need to understand molecular alterations associated with genomic instability and the specific genes that can drive DNA repair defects. In order to comprehensively identify tumors with HR defects, we utilized a genomic scar HRD score. This score allows us to detect genomic lesions left by HRD and identify tumors with HRD regardless of the genetic event which caused the HRD (Telli et al., 2016). HRD scores calculated across tumors from The Cancer Genome Atlas (TCGA) revealed numerous molecular alterations associated with HRD. Surprisingly, approximately 75% of tumors that scored positive for HRD exhibited no known molecular HRD driver. Using an integrated network-based approach, we identified nearly 100 novel candidate HRD drivers, with a particular enrichment for genes involved in RNA processing. Candidate HRD drivers had over a 90% experimental validation rate. Mechanistic studies indicated that that the RNA binding proteins we identified may influence HR either by modulating expression of canonical DNA repair genes, or by directly acting directly at sites of DNA damage.

## RESULTS

### HRD scores vary across tumor types and patient demographics

HRD leaves a quantifiable genomic scar allowing the calculation of an HRD score, defined as the combination of three measures of genomic instability: loss of heterozygosity (LOH), telomeric allelic imbalance (TAI), and large-scale transitions (LST) (Fig. 1A) (Telli et al., 2016). We determined the HRD score across tumors from all patients within TCGA, and found high HRD scores for basal-like breast cancer and ovarian cancer, both of which are known to have high levels of HRD (Couch et al., 2014; Konstantinopoulos et al., 2015). Luminal androgen receptor, and luminal A and luminal B breast cancers had low HRD scores (Fig. S1A-B). In addition to basal-like/ovarian cancer, numerous other cancer types not typically associated with HRD exhibited high HRD scores, including lung squamous carcinoma (LUSC), bladder cancer (BLCA), and gastric cancer (STAD) (Fig. 1B). Consistent with early onset of basal/triple-negative breast cancers (TNBC), we found that in tumors from basal breast patients, HRD score was negatively associated with patient age. There were similar trends observed in lung adenocarcinoma (LUAD), head and neck squamous carcinoma (HNSC), and mesothelioma (MESO). Nonetheless, when quantified across all patients, HRD score generally showed a positive relationship with age (P = 5.3×10^-7^, Fig. 1B, Fig. S1C). When compared across all cancer types, tumors from male patients tended to have higher HRD scores than female patients (P = 0.04, Fig. 1B, Fig. S1D), and tumors from Asian patients had statistically higher HRD scores than other patients (P = 4.3×10^-3^, Fig. 1B, Fig. S1E-F).

**Figure 1.**
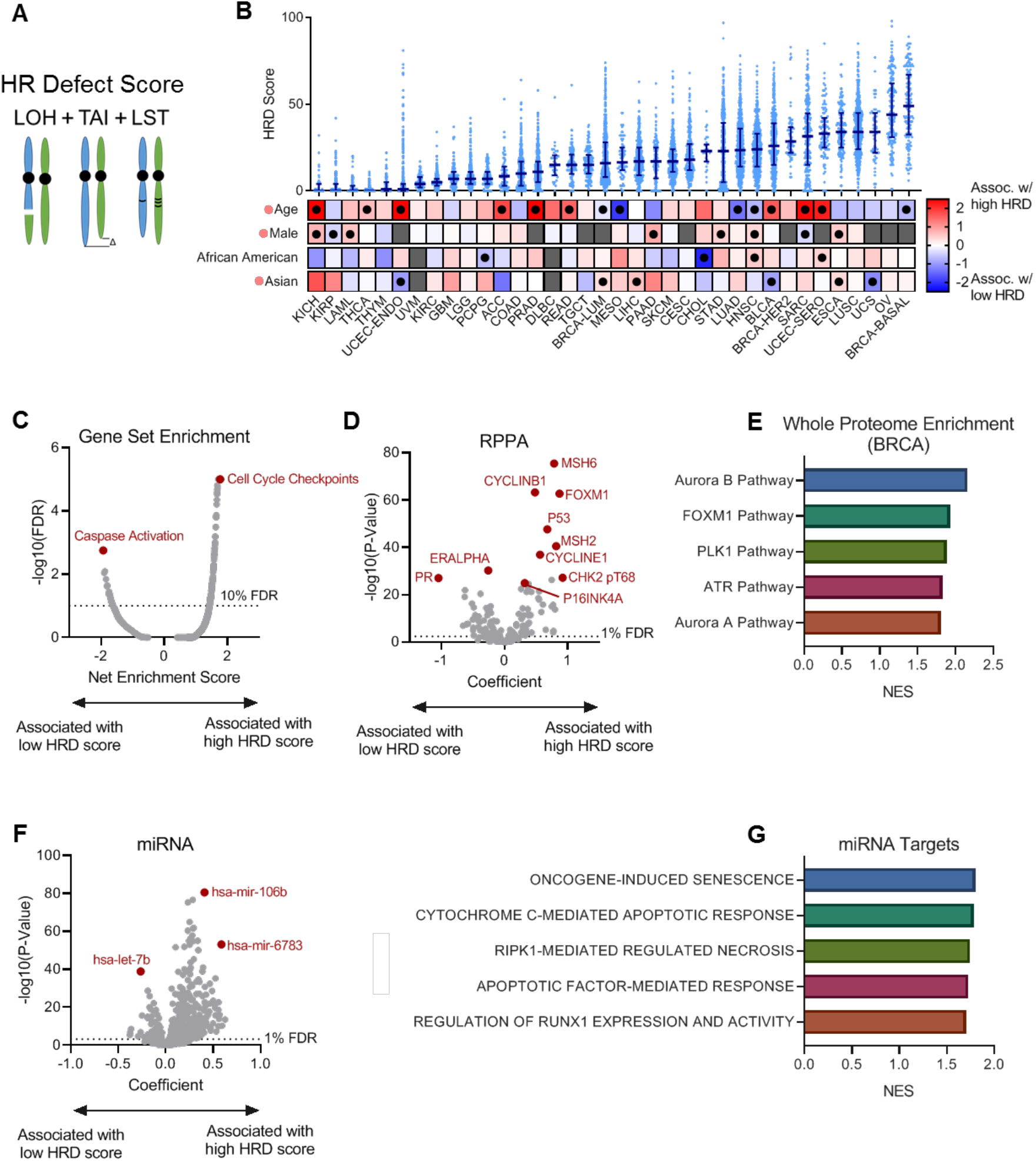
Pan-cancer analysis of HR defects. **(A)** Schematic describing the HR defect score, defined as the sum of scores for loss of heterozygosity (LOH), telomere allelic imbalance (TAI), and large scale state transitions (LST). **(B)** HRD scores across tumor types (top), and associations with various demographic features (bottom). HRD score is plotted as median value with error bars representing interquartile range. Demographic features scale from red (positive association with HRD score) to blue (negative association with HRD score). Black dots represent significant relationships within a given cancer type. Red dots next to demographic categories represent a significant positive association across all cancer types. **(C)** Gene set enrichment based on relationship with HRD score. A generalized linear mixed model was used to determine the association between RNAseq-derived gene expression levels and HRD scores, taking tumor type as a random effect. The resulting coefficients were used for gene set enrichment analysis. **(D)** Association between HRD score and protein levels. A generalized linear mixed model was used to determine the association between reverse phase protein array-derived protein levels and HRD scores, taking tumor type as a random effect. FDR was determined using the Benjamini– Hochberg procedure. **(E)** Pathway enrichment determined from whole-proteome mass spectrometry in breast cancer. Spearman correlation coefficients were determined between proteins and HRD scores. The resulting correlation coefficients were used for gene set enrichment analysis. The five most significantly enriched pathways are shown. NES = normalized enrichment score. **(F)** Association between HRD score and microRNA levels. A generalized linear mixed model was used to determine the association between microRNA levels and HRD scores, taking tumor type as a random effect. FDR was determined by the method of Storey. **(G)** Gene set enrichment of microRNA target genes. For each gene, a score was quantified as the sum of all coefficients for significant (FDR<1%) microRNAs that could target that gene. The resulting list of scores was used for GSEA. The five most significantly enriched pathways are shown, demonstrating pathways predicted to be suppressed by miRNAs associated with HRD score. NES = normalized enrichment score. See also Figure S1.

### HRD scores are associated with suppression of cell death pathways and activation of DNA damage response checkpoints

To begin to define the molecular changes associated with HRD across cancer types, we performed gene set enrichment analysis (GSEA) (Subramanian et al., 2005) and found that HRD scores were associated with increased expression of cell cycle checkpoint genes and decreased expression of genes associated with caspase activation (Fig. 1C). As shown in Fig. 1D, protein-level analysis using reverse phase protein array (RPPA) data indicated HRD score was positively associated with numerous cell cycle regulators, including the DNA damage checkpoint marker phospho-CHK2, as well as MSH6 previously implicated non-homologous end-joining outside of its canonical role in DNA mismatch repair (Shahi et al., 2011). We confirmed the association of HRD score with an increased expression of cell cycle checkpoint proteins in breast tumors using an orthogonal whole-proteome mass spectrometry-based dataset (Fig. 1E) (Mertins et al., 2016). Analysis of microRNAs (miRNAs) again identified numerous miRNAs associated with HRD score (Fig. 1F). We desired to assess which pathways these miRNAs might modulate, but the analysis was complicated: each miRNA can target multiple genes and each gene can be targeted by multiple miRNAs. To address this complication, we calculated a gene-wise miRNA suppression score for each gene, defined as the sum of the miRNA coefficients predicted to target each specific gene. Final miRNA scores for all genes were used for GSEA. We found that miRNAs associated with high HRD scores preferentially suppressed oncogene-induced senescence and apoptotic pathways (Fig. 1G). Taken together, these results indicate that HRD positive tumors tend to have suppressed tumor suppressor pathways and activated DNA damage checkpoint pathways.

### The majority of HRD are of unknown aetiology

In order to bifurcate tumors into HRD positive and HRD negative groups, we took a multi-step approach. We began by analyzing tumors of patients with known deleterious *BRCA1* or *BRCA2* germline mutations as a gold standard for HRD positivity. We focused this analysis across breast, ovarian, pancreatic, and prostate cancer where *BRCA1* or *BRCA2* germline mutations are known to promote tumorigenesis. As indicated by the receiver-operator characteristic curve in Figure 2A, HRD score was highly accurate at recovering tumors with deleterious *BRCA1* or *BRCA2* germline mutations, demonstrated by an area under the curve (AUC) value of 0.83. Using this data, we determined an optimal HRD threshold score of 32, which identified numerous HRD positive tumors across an array of different cancer types (Fig. 2B). However, an analysis of the genetic events within these tumors indicated that only a small minority (9.7%) displayed alterations (mutation or methylation) in *BRCA1*/*BRCA2.* Therefore, we expanded the analysis of genetic events to include HR associated genes from a larger, annotated list (Lord and Ashworth, 2016) but could still only identify potential drivers for roughly 25% of HRD tumors (Fig. 2C).

**Figure 2.**
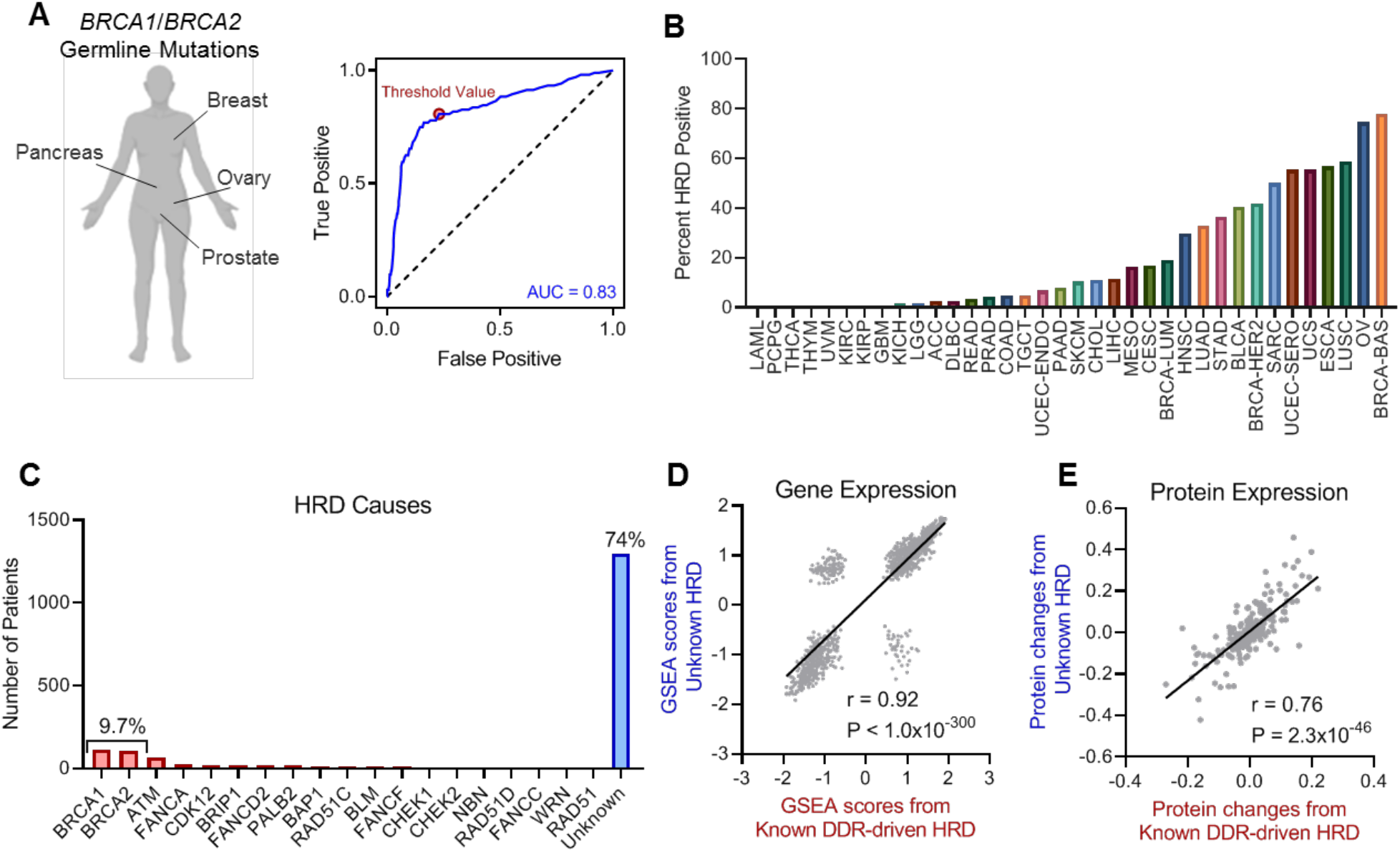
Most HR deficiencies are of unknown aetiology. **(A)** Receiver operator characteristic curve (ROC) (blue) demonstrating the ability of the HRD score to predict germline mutations in *BRCA1* or *BRCA2* in the indicated tumor types. AUC is defined as the area under ROC curve. The dotted line represents the expectation due to random assignment. The red dot indicates the calculated optimal threshold value to separate HRD positive (HRD^+^) and HR competent tumors. **(B)** Percentage of HRD positive tumors by cancer type. **(C)** Percentage of HRD positive tumors caused by *BRCA1*, *BRCA2*, other canonical DDR genes, and those of unknown aetiology. **(D)** Correlation of gene set enrichment analysis performed on gene expression changes between DDR-driven HRD positive tumors and HR competent tumors (plotted on x-axis), or HRD positive tumors of unknown aetiology and HR competent tumors (plotted on y-axis). Spearman correlation coefficient. **(E)** Correlation of change in RPPA-derived protein levels between DDR-driven HRD positive tumors and HR competent tumors (plotted on x-axis), or HRD positive tumors of unknown aetiology and HR competent tumors (plotted on y-axis). Spearman correlation coefficient.

The large fraction of HRD positive tumors with causes that could not be attributed to known drivers of HRD may represent false positives, or alternatively, could indicate that the majority of drivers of HRD in patients with cancer are unknown. Based on the strong molecular alterations we observed to be associated with HRD (Fig. 1C-G), we hypothesized that if the HRD of unknown origin were due to true HRD, they would exhibit similar molecular changes to those caused by known drivers such as *BRCA1*/*BRCA2*. Analysis of GSEA scores from gene expression data were correlated for two groups: 1) DDR-driven HRD-positive tumors vs. HR-competent tumors, and 2) HRD tumors with unknown causes vs. HR-competent tumors. Both groups demonstrated remarkably concordant changes in gene expression (Fig. 2D). Likewise, correlation analysis performed at the protein level using RPPA data also revealed a robust correlation between protein alterations in HRD positive tumors with known DDR gene alterations and those of unknown origin (Fig. 2E). These data indicate that a large fraction of HRD positive tumors are driven by unknown causes.

### Network-based discovery of novel drivers of HRD

Across all tumor types, approximately 75% of HRD positive tumors had an intact complement of known HR-related genes, indicating that the majority of HRD drivers were of unknown aetiology (Fig. 3A). To define the causes of HRD in these tumors, we developed a network-based algorithm for identifying novel drivers of HRD (Fig. 3B). We began with a list of verified inducers of HRD. Then we identified genetic events in the genes encoding these inducers in patient tumors. Tumors were considered to have a genetic event if they had either: 1) mutations with high variant allele frequency (VAF), or 2) a methylation event that corresponded with downregulation of the methylated gene. VAF was used for assessment because if a mutation is driving HRD/tumorigenesis, then it should occur in the majority of tumor cells. Next, we assessed whether a genetic event was associated with an increased HRD score based on cancer type. Candidate genetic events that may drive HRD in an individual tumor were assigned based on the degree to which that event increased the HRD score, the number of tumors in which it occurred, and the VAF of the mutation in a given tumor. After assigning the genetic events that may cause HRD to an individual tumor, we hypothesized that proteins that interact with these drivers would also be more likely to cause HRD. Therefore, we used protein-protein interaction (PPI) networks to expand the list of candidate HRD drivers. This prediction algorithm was iterated until convergence, when it could no longer identify additional putative novel HRD driver candidates.

**Figure 3.**
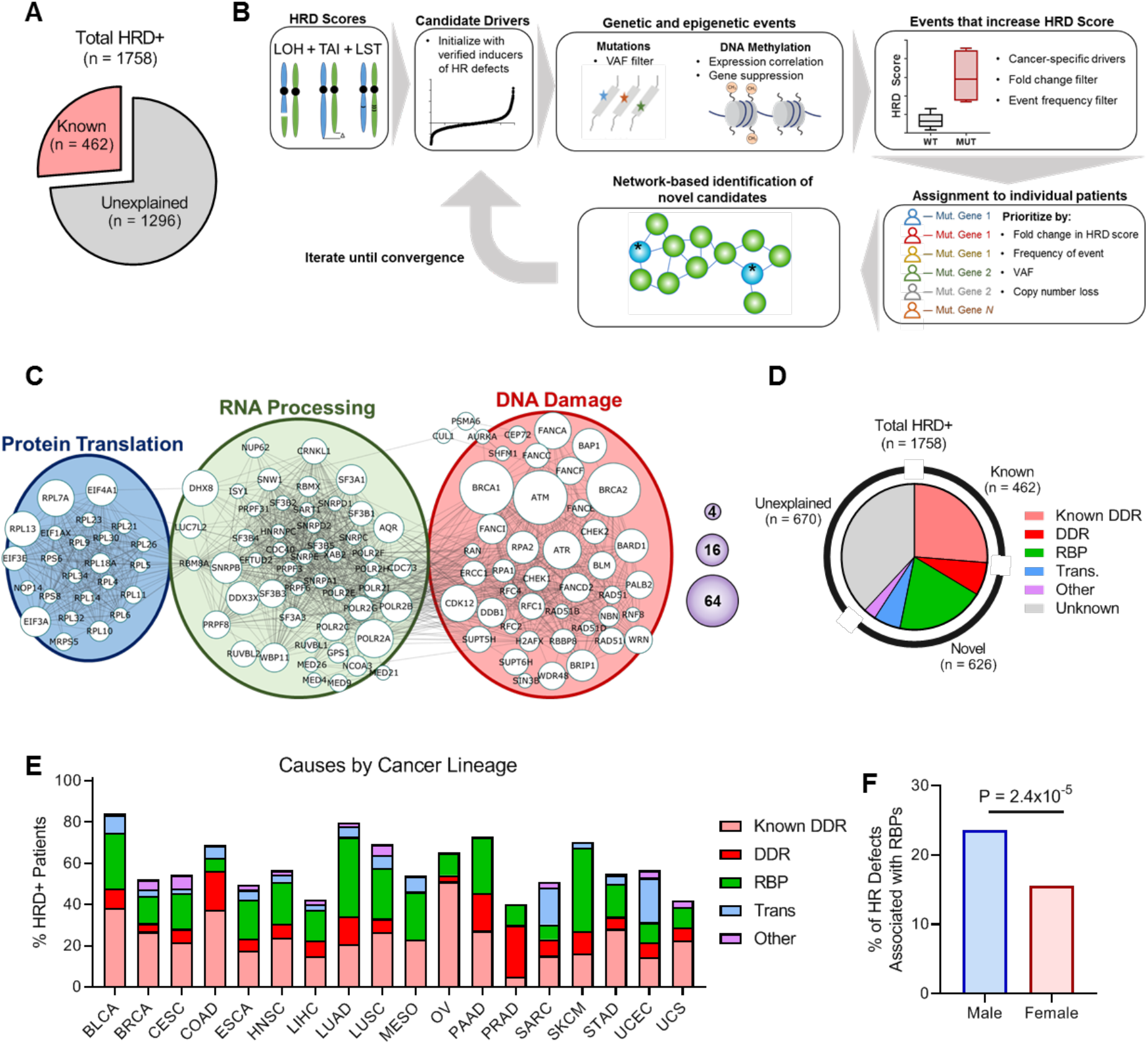
Discovery of novel drivers of HRD. **(A)** Pie chart showing the fraction of HRD positive (HRD+) tumors with known drivers of HRD and those with unknown drivers of HRD. **(B)** Schematic of the network-driven pipeline used to discover novel drivers of HRD. **(C)** Resulting protein network modules of identified drivers of HRD in tumors from patients with cancer. The size of nodes represents the number of patients with HRD attributed to a given gene, corresponding to the scale shown in the purple spheres. **(D)** Pie chart showing the fractions of previously identified HRD drivers (light red) and newly identified putative drivers. **(E)** Distribution of HRD driver ontologies by cancer type. **(F)** Percentage of HR defects driven by RBPs in tumors from male and female patients with cancer. Fisher’s exact test. See also Figure S2-S3 and Table S1.

The final network of predicted HRD drivers (Fig. 3C) could be grouped into three, large protein modules. As anticipated, the largest of these was a DNA damage module, followed by an RNA binding protein (RBP) module, and a smaller protein translation module. Only a small fraction of proteins (< 3%) was not strongly associated with any one of these modules. In total, we identified novel HRD drivers for 626 of 1296 patient tumors displaying HRD but lacking a discernable alteration in a previously defined HR pathway (unexplained HRD), with the largest fraction consisting of RBP genes (Fig. 3D). Identified HRD causes and ontology annotations for each tumor are given in Table S1. Tumors that we failed to identify a putative driver for exhibited significantly lower (P = 5×10^-11^) HRD scores, suggesting these samples may be enriched for false positives. The relative proportion of various HRD drivers varied across tumor types. Ovarian, bladder, and colorectal cancers showed the largest fractions driven by canonical DDR genes, whereas melanoma, lung adenocarcinoma, and pancreatic cancer, showed the largest fractions driven by RBP genes (Fig. 3E). Notably, tumors from men were more likely to have HRD driven by RBP mutations (Fig. 3F). The majority of patients with bladder cancer in the TCGA cohort received cisplatin or similar chemotherapies, and should respond favorably if the tumor is HRD. We found that all identified causes of HRD were associated with good prognosis in bladder cancer, indicating the identified HRD drivers likely contribute to HRD and thus chemosensitivity (Fig. S2).When we analyzed the remaining HRD positive tumors with unexplained HRD, we identified several miRNAs that could suppress the expression of HRD drivers that were up-regulated in HRD positive sarcoma tumors (Fig. S3).

### Functional validation of novel drivers of HRD

We next sought to validate that the novel putative drivers of HRD identified from patient tumors were functionally important in HR and not simply bystander events. To begin to validate the functional relevance of candidate genes, we began by utilizing the DR-GFP reporter assay. In this assay, expression of an eGFP variant can only be restored if it is accurately repaired by HR following cleavage with I-SceI (Pierce et al., 1999). Pladienolide B, which inhibits the core spliceosome RBP SF3B1, significantly inhibited HR at concentrations as low as 1 nM in U2OS cells, suppressing HR comparable to inhibition of the Mre11-Rad50-Nbs1 complex critical for HR (Ciccia and Elledge, 2010) with 100 μM Mirin (Fig. S4A). For more specific analysis, we utilized two independent siRNAs for 43 of the potential HRD mediators we identified, and found that 95% induced HR defects (Fig. 4A). The two genes that failed to induce HR defects, *CEP72* and *JMJD6*, were both classified as “other.” The genes in this category were only weakly linked to any module, further indicating that our network-based approach increased the robustness of our ability to identify strong candidate drivers. To exclude the potential that reduced HR function is merely an artifact from arresting cell cycle, we analyzed the ability of cells to form irradiation (IR)-induced Rad51 foci specifically in cycling cells following suppression of candidate RBPs, revealing highly concordant results (Fig. S4B-C). A complete summary of the results of functional assays are given in Table S2. As depletion of RBP proteins may not be functionally equivalent to the effects of missense mutations, we engineered vectors expressing RBPs with mutations identified from HRD tumors. Cells were transiently transfected with RFP-tagged mutant proteins or wild-type controls, allowing for analysis of HR function in cells expressing the desired constructs. After we profiled 29 mutations derived from patient tumors across 10 genes, we found that 28/29 inhibited HR function (Fig. 4B).

**Figure 4.**
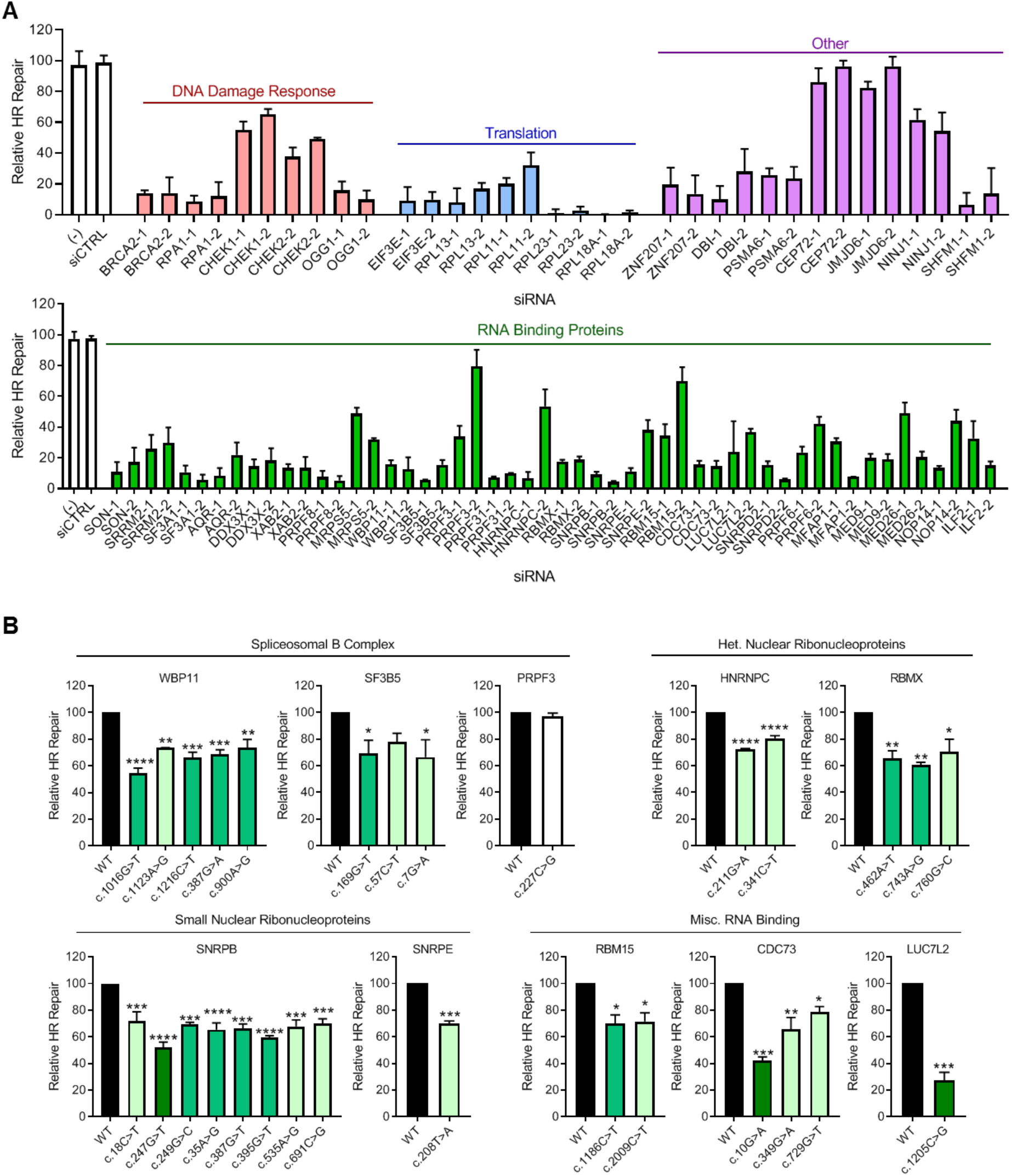
Validation of novel drivers of HR defects using flow cytometry-based DR-GFP reporter assay. **(A)** HR repair was measured using a DR-GFP assay and flow cytometry to detect GFP in cell lines following siRNA-mediated knockdown of each of the indicated genes. Relative HR repair is defined relative to the percentage of GFP positive cells of untransfected and siCTRL-transfected controls. N = 3 per siRNA, 2 siRNAs per gene. Mean ± s.e.m. **(B)** The relative amount of HR repair was measured as in A. Cells were transfected with RFP-tagged wild-type (WT) or mutant protein-expressing constructs 24 hours prior to transfection with an I-SceI expression plasmid. HR repair was measured in RFP+ cells, and defined as mutant relative to WT overexpression. N = 3 per condition. Mean ± s.e.m. ANOVA with Dunnett’s post-hoc test. *p<.05, **p<0.01, ***p<0.001, ****p<1×10^-4^

To further validate the role of RBPs in HR, we treated TNBC MDA-MB-231 cells with pladienolide B, and found that it efficiently inhibited the formation of IR-induced Rad51 foci in cycling cells (Fig. 5A-B). As PARP inhibitors are known to preferentially kill HRD cells, we hypothesized that the RNA spliceosome inhibitor pladienolide B would sensitize these cells to PARP inhibitors. Indeed, we found pladienolide B and the PARP inhibitor BMN-673 demonstrated synergy in two TNBC cell lines, MDA-MB-231 (Fig. 5C) and BT-549 (Fig. 5D). For more specific analysis, we repeated the IR induced Rad51 foci assay following siRNA-mediated depletion of DDX3X and AQR, two of the most common mutant RBPs in TNBC. We found that both siDDX3X and siAQR both inhibited foci formation across 4 TNBC cell lines (Fig. 5E). We further validated that suppression of AQR likewise suppressed HR in ovarian cancer cells, where it was also predicted to be associated with HR function (Fig. S4D). As HR deficient tumors may be therapeutically targeted with PARP inhibitors, we next tested the effects of PARP inhibitors in MDA-MB-231 cells stably expressing shDDX3X, shSF3B3, or shBRCA2. We found that depletion of either of the RBPs DDX3X or SF3B3 increased cell sensitivity to the PARP inhibitors BMN-673 (Fig. 5F) and AZD2281 (Fig. 5G) as well as if not better than depletion of BRCA2. Long-term clonogenic assays performed in the presence of BMN-673 confirmed this increased sensitivity to PARP inhibition (Fig. 5H). Together, these results indicate that the novel HRD drivers we identified are likely to be functionally relevant for HR repair across multiple cell lines and assays.

**Figure 5.**
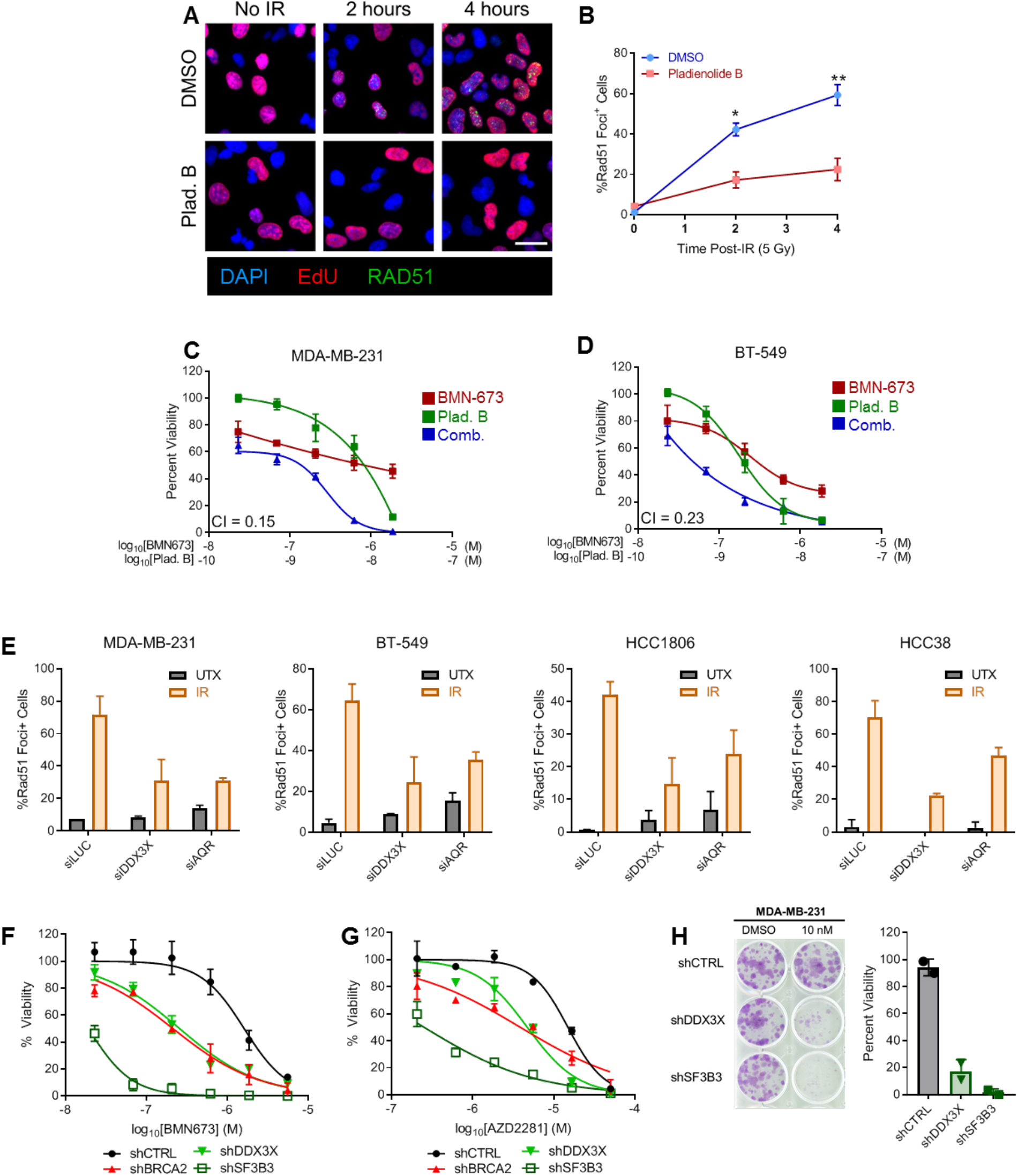
Control of HR by RNA binding proteins in breast cancer. **(A)** Images of cells treated with either 10 nM of the splicing inhibitor pladienolide B or a DMSO vehicle control, 24 hours prior to irradiation (5 Gy). Images show Rad51 foci indicative of HR repair (green), as well as proliferating (S phase) cells with EdU (red), and DAPI nuclear stain (blue). **(B)** Quantification of percentage of cycling (EdU+) cells that exhibit Rad51 foci. N = 3. Student t-test. *p<.05, **p<0.01 **(C-D)** Viability of TNBC cells following a 5-day incubation with the splicing inhibitor pladienolide B (concentration indicated on top x-axis), PARP inhibitor BMN673 (concentration indicated on bottom x-axis), or a combination thereof, relative to a DMSO vehicle control for MDA-MB-231 cells (C) and BT-549 cells (D). Mean ± s.e.m. C.I.; Chou-Talalay combination index. N = 2. **(E)** Quantification of IR-induced Rad51 foci in cycling (EdU^+^) TNBC cell lines following siRNA-mediated depletion of the indicated RBPs. N = 2 per cell line. **(F)** Viability of individual MDA-MB-231 cell lines with stable single knockdowns of RBPs DDX3X and SF3B3, as well as HR protein BRCA2, following a 5-day treatment with BMN673. **(G)** As in F but following a 5-day treatment with AZD2281. **(H)** Clonogenic assay with MDA-MB-231 cells with stable knockdown of RBPs DDX3X and SF3B3, following a two-week treatment with BMN673. N = 2, points represent independent biological replicates. See also Figure S4.

### Induction of HR defects may occur by modulation of DDR genes as well as independent pathways

Next, we sought to understand how the newly identified novel drivers of HRD might influence HR repair. We hypothesized that loss of RBPs could interfere with DDR gene expression, by either decreasing mRNA stability or by interfering with splicing, either of which could result in hindered protein function. To test whether RBPs were modulating DDR genes, we performed multiple experimental and computational analyses. First, RNAseq analysis following siRNA-mediated depletion of 17 RBPs in three cell lines to identify differential DDR gene expression relative to either siCTRL, or siBRCA1/siBRCA2. Next, TCGA patient tumors were analyzed to detect decreased DDR gene expression relative to HR competent tumors or tumors with HRD caused by DDR genes. Finally, the same comparisons were made using TCGA alternative splicing analysis rather than gene expression levels. All comparisons were made to both siCTRL/HR competent samples and siDDR/HRD caused by DDR, as we had found that HRD itself can cause transcriptional rewiring (Fig. 1C, 2D). The integration of these results is shown in Fig. 6A, with specific comparisons shown in Fig. S5. We identified DDR genes that were either suppressed or alternatively spliced for 55% (26 out of 47) of the RBPs that are candidates for affecting HR. The largest influence on gene expression was seen for members of the mediator complex (75% of genes), followed by core spliceosome members (61% of genes), with less influence seen by other RNA binding proteins (36% of genes). We validated 10 proteins identified to be modulated at the gene expression level by 3 RBPs at the protein level by western blot, and found all candidates showed lower levels of protein as expected (Fig. S5G-I).

**Figure 6.**
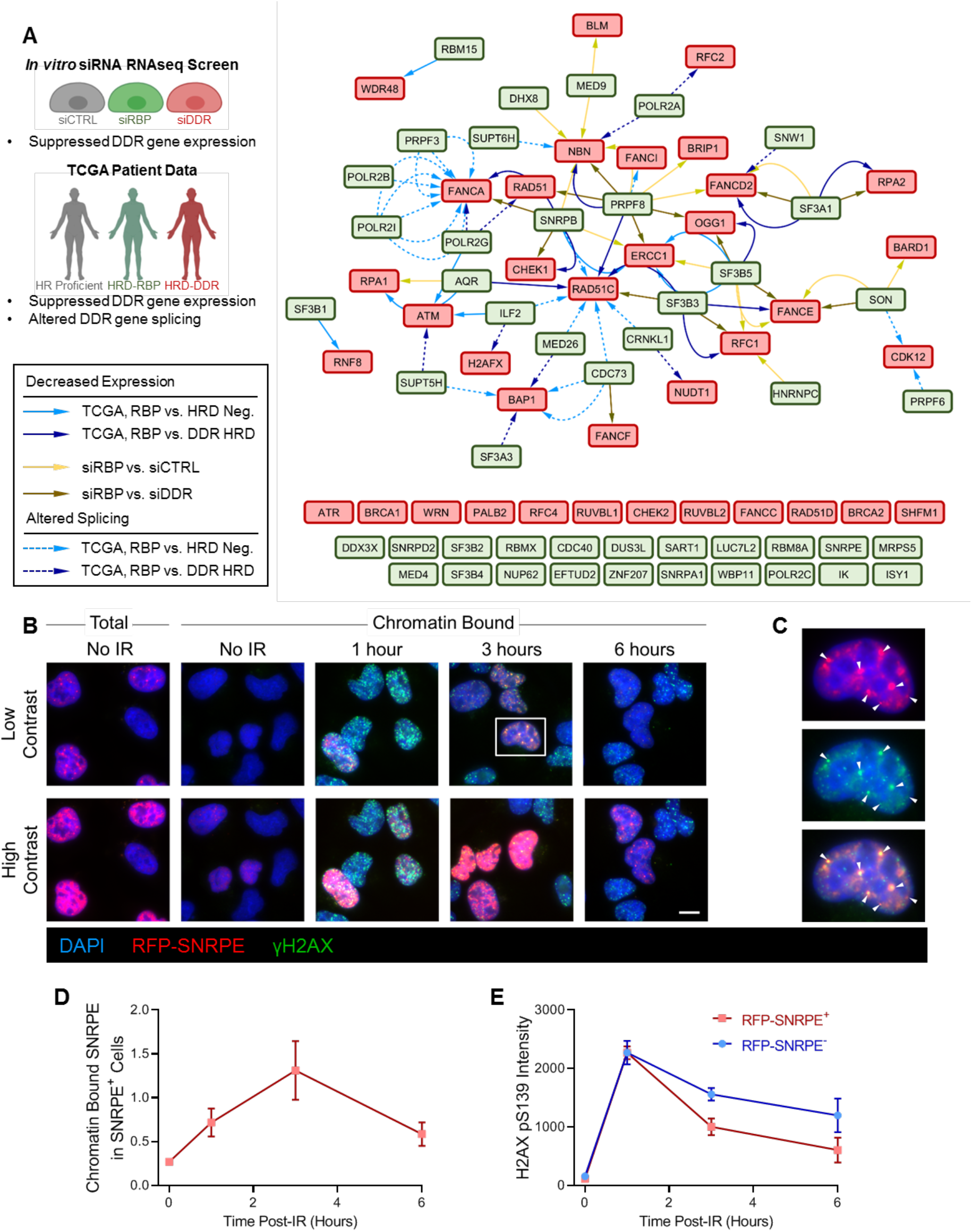
RNA binding proteins control HR repair by multiple mechanisms. **(A)** RBPs that can modulate DDR gene expression were identified through three approaches: (1) *in vitro* RNA-seq screening following siRNA-mediated knockdown of selected (N = 17, Table S2) pipeline-identified RBPs compared to either control siRNA (siCTRL) or siBRCA1 and siBRCA2 (siDDR) in isogenic cell lines; (2) identification of down-regulated DDR genes expression levels in TCGA tumors with HRD driven by RBP loss compared to HR competent tumors or tumors with HRD driven by DDR loss; and (3) identification of alternatively spliced DDR genes in TCGA patient tumors with HRD driven by RBP loss compared to HR competent tumors or tumors with HRD driven by DDR loss. The network nodes indicate RBPs (green) identified to modulate DDR genes (red), with edges indicating how the modulation was identified. Genes listed across the bottom were not identified as being modulated by RBPs. The key for the network diagram, with solid arrows representing decreased expression and dotted arrows representing altered splicing, is on the left. **(B)** IR-induced foci formation showing merged images of co-staining of RFP-SNRPE (red) and γH2AX (green). Total and chromatin-bound proteins prior to irradiation and chromatin-bound proteins at 1, 3, and 6 hours following irradiation (5 Gy). Scale bar = 10 μm. **(C)** Magnified image of boxed cell indicated in (B) showing single channel SNRPE (top) and γH2AX (middle), and the merged image (bottom). Arrows mark foci with co-localization of RFP-SNRPE and γH2AX. **(D)** Quantification of fluorescence signal due to chromatin-bound RFP-SNRPE in SNRPE^+^ cells prior to and at 1, 3, and 6 hours following irradiation. Intensity is reported relative to the median total intensity in cells not subjected to extraction. **(E)** Quantification of the fluorescence signal due to γH2AX prior to and at 1, 3, and 6 hours following irradiation in RFP-SNRPE^+^ and RFP-SNRPE^-^ cells. See also Figure S5-S6.

For those RBPs that did not appear to be directly modulating expression of DDR genes, we hypothesized that they might act directly at sites of DNA damage. To evaluate whether RBPs might act at sites of damage, we tested whether the RBP SNRPE formed foci in response to IR-induced damage. Although at baseline SNRPE is largely nuclear, it is not tightly chromatin bound and most can be extracted (Fig. 6B). Within 1 hour after cells were irradiated, SNRPE became increasingly chromatin bound, and co-localized with the DNA DSB marker γH2AX (Fig. 6C-D). Furthermore, cells over-expressing SNRPE demonstrated quicker DSB repair as quantified by quicker loss of γH2AX foci (Fig. 6E).

Based on the ability of RBPs to modulate expression of DDR genes, we hypothesized that a similar paradigm may apply to mutations in genes regulating translation by altering protein levels of DDR proteins. However, our analysis of TCGA RPPA data for DDR proteins revealed no significant relationships between mutations in translation genes, such as E4F1, and decreased DDR protein expression (Fig. S6A-B). Nonetheless, as observed for SNRPE, we were able to detect IR-induced E4F1 foci (Fig. S6C), suggesting that E4F1 might have a functional role at sites of DNA damage.

### Novel HR drivers generalize to independent cohorts and are associated with cancer risk

While complete validation of all novel HRD drivers would be time prohibitive, to evaluate whether the novel HR drivers we uncovered in the TCGA data might be more broadly applicable to patient tumors in general, we analyzed additional patient cohorts. Interrogation of the International Cancer Genome Consortium (ICGC) breast cancer patient cohort validated that mutations from all identified ontologies were significantly associated with higher HRD scores (Fig. 7A). Furthermore, RBP mutants accounted for a similar fraction of HRD tumors as that observed in the TCGA cohort (Fig. 7B, Fig. 3E). The association of HRD score with candidate drivers was maintained when analyzing TNBC tumor alone (Fig. 7C).

**Figure 7.**
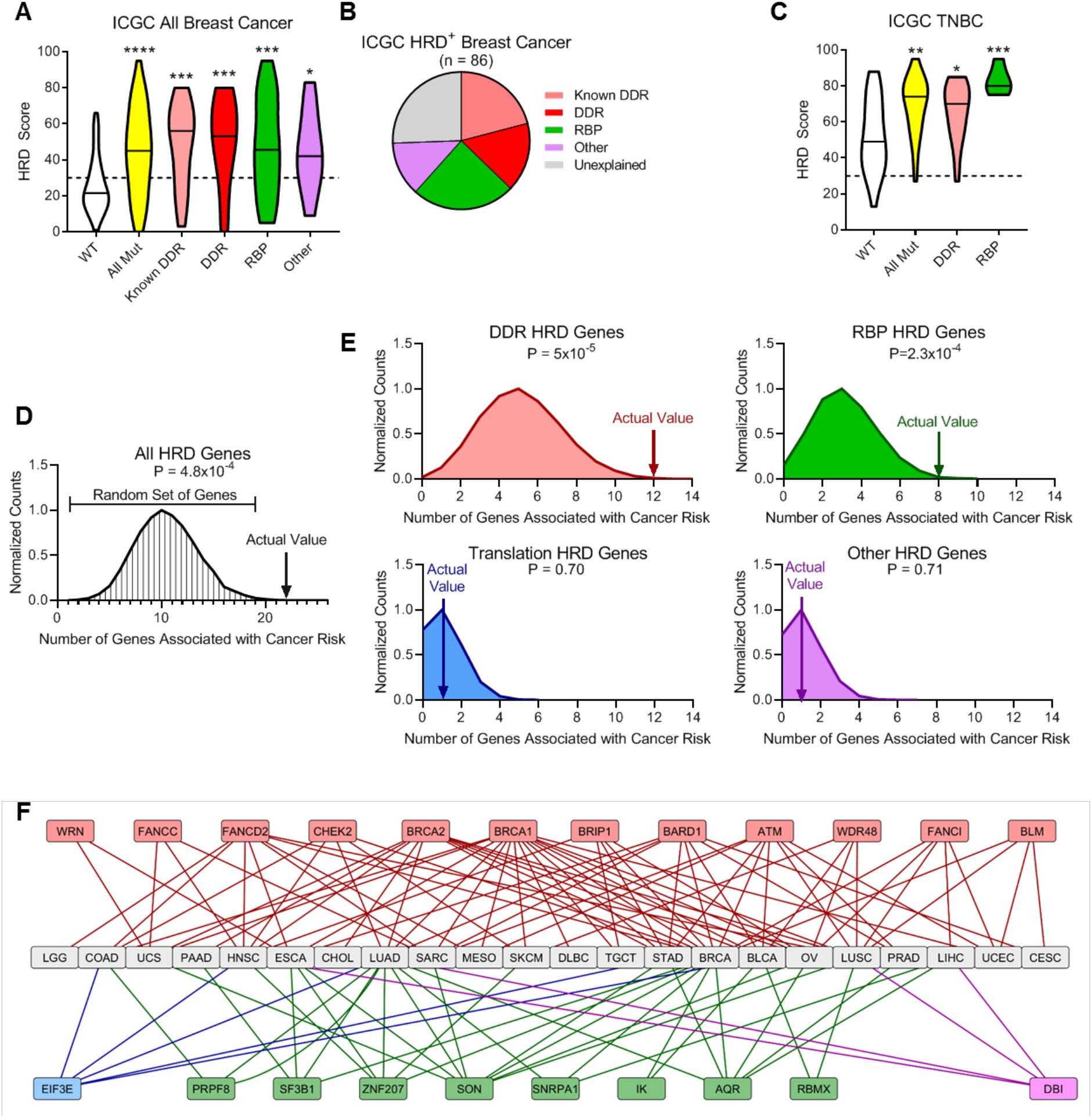
External validation of the involvement of RNA binding proteins in HR repair. **(A)** HRD scores were determined for an independent cohort of tumors from patients with breast cancer from ICGC. Tumors were classified based on their candidate drivers into either Known DDR (pink), novel DDR (red), RBP (green) or other (purple) genes. Kruskal-Wallis with Dunn’s posthoc comparing each group to WT. **(B)** Pie chart showing the relative proportions of proposed causes of HRD defects in patient tumors from the ICGC cohort. **(C)** HRD scores as in (A), but only showing tumors form TNBC patients. Kruskal-Wallis with Dunn’s posthoc. **(D)** Identified novel drivers of HR defects are enriched for genes associated with cancer risk from genome-wide association studies. The graph shows the distribution of number of genes associated with cancer risk using a random set of genes of equal size. The arrow indicates the actual observed number of genes associated with cancer risk. Empirical P value. **(E)** Enrichment in genes associated with cancer risk as described in (D), but for individual gene ontologies. **(F)** Network showing genes identified from the GWAS studies associated with cancer risk as well as the cancer types where mutations in those genes were identified as causing an HR defect. DDR genes (red), RBP genes (green), translation genes (blue), and other (purple). *p<.05, **p<0.01, ***p<0.001, ****p<1×10^-4^

As loss of function in genes associated with HR is associated with increased propensity for cancer development, the genes we identified as candidate drivers of HRD should be enriched in genes associated with cancer risk in genome-wide association studies (GWAS). Therefore, we assembled all genes associated with cancer risk from GWAS DB (Li et al., 2016) and looked for the HRD drivers identified through our pipeline. Overall, we found that our identified HRD drivers were significantly enriched for genes associated with cancer risk (Fig. 7D). Analysis of individual gene ontology indicated that this enrichment was largely driven by DDR and RBP genes (Fig. 7E). Mutations in genes associated with cancer risk were detected across numerous cancer types (Fig. 7F). These mutations, when occurring in the germline, could provide vital information for genetic counseling to improve cancer screening/prevention.

## DISCUSSION

Our analysis of genomic scars indicative of HRD across tumors in the TCGA indicated that only about 25% of HRD could be attributed to alterations in known drivers of HRD. The remaining 75% of HR-deficient tumors had no identifiable defects in known DDR genes. However, these tumors displayed gene and protein expression changes consistent with HRD caused by aberrations in DDR genes known to cause HRD, including activation of cell cycle checkpoints and suppression of senescence/apoptosis pathways. Suppression of senescence/apoptosis pathways may be critical for HRD tumor cells to circumvent tumor-suppressive DDR checkpoints (Bartkova et al., 2006), thus enabling genomically unstable cells to continue proliferating and acquire additional mutations. This transcriptional re-wiring may explain how tumor cells continue to proliferate in absence of *BRCA1*/*BRCA2*, whereas depletion of these genes in non-malignant cells often reduces cellular fitness. Using a networks-based approach, we more than doubled (from 462 to 1088) the number of tumors in TCGA with an attributable driver of HRD. Among the novel drivers, we found a particular enrichment for aberrations in genes encoding RNA binding proteins, which represented over half of newly identified drivers of HRD.

The potential role of RBPs in controlling HR is consistent with limited, previously published reports. For example, genome-wide siRNA depletion screens designed to evaluate HR function identified RBPs, namely *RBMX* (Adamson et al., 2012) and *CDC73* (Herr et al., 2015), as potential drivers of HRD. An orthogonal genome-wide screen for PARP inhibitor sensitivity using CRISPR-mediated deletion also recovered genes consistent with our results, including *SF3B3* and *SF3B5* (Zimmermann et al., 2018). However, *in vitro* screening approaches do not necessarily correspond to *bona fide* drivers of HRD observed in patient tumors. Genes identified as critical for HR through loss of function screens might be essential genes, meaning loss is incompatible with cell viability, or may simply not be mutated at appreciable frequencies in human populations. Additionally, the deletion/depletion of genes may not reflect the phenotypes observed when those same genes are mutated. The disparity between *in vitro* data and observations in patient tumors is best highlighted by the two aforementioned siRNA screens that identified 6,137 (Adamson et al., 2012) and 10,050 (Herr et al., 2015) genes that reduced HR function more than the average reduction observed following loss of canonical HR/BRCAness genes (Lord and Ashworth, 2016). In contrast, our study indicated that only 1.58% of the 6,137 genes and 1.04% of the 10,050 genes may be relevant in tumors from patients with cancer.

The genomic scar HRD score calculation and corresponding analyses performed in our study are subject to several limitations. The primary limitations are centered around the accuracy of the HRD score itself, as well as the chosen threshold for HRD positive and negative tumors. Although the combination of three different measures of genomic instability offers an improved signal over any single metric, it may still not capture all HRD tumors (Telli et al., 2016). Further, we assumed a constant threshold value for HRD positivity across all tumor types, but the validity of this assumption is unclear. At the molecular level, we observed consistent changes between HRD driven by canonical drivers and those with originally unidentified aetiology, suggesting that HRD score reflects loss of HR function in both contexts and that the HRD positive tumors of unidentified aetiology are not generally false positives. However, at the single tumor level the HRD score is still subject to false positives/negatives. For example, at the optimal threshold identified for bifurcation into HRD positive and negative tumors, roughly 20% of *BRCA1*/*BRCA2* germline mutations were classified as existing in HR competent tumors. The observed false negative germline *BRCA1*/*BRCA2* mutations could represent actual false negatives, or may represent non-deleterious variants of *BRCA1*/*BRCA2*. By enforcing occurrence of genetic events in multiple tumors, we were largely able to avoid the effects of sporadic false positives, as evidenced by the over 90% experimental validation rate.

The findings presented in this study warrant follow up studies to further elucidate the mechanisms underlying how the newly identified drivers of HRD control HR repair. Previous reports have documented the role of RBPs in controlling expression of DDR factors, for example, RBMX has been shown to be required for BRCA2 expression (Adamson et al., 2012). We likewise detected that 26 and 47 RBPs analyzed may suppress expression of canonical HR genes. Consistent with our observation that ILF2 can modulate multiple DDR genes, ILF2 overexpression facilitates expression of DDR genes leading to resistance to DNA damaging agents in 1q21-amplified multiple myeloma (Marchesini et al., 2017). Following DNA damage, the spliceosome has also been shown to undergo rapid mobilization resulting in alternative splicing that may facilitate repair of DNA lesions (Tresini et al., 2015). Although analysis of Rad51 foci formation indicates RBPs largely induce HR defects at or before Rad51 loading, future studies to elucidate precisely which step(s) of HR putative drivers of HR defects are responsible for are warranted. Alternatively, RBPs may not interact directly with HR proteins but interfere with generation of RNA species required for HR repair. For instance, recent work indicated that DICER and DROSHA RNA products are involved in activation of the DDR, and that DDR foci can be abrogated by RNase A-mediate RNA degradation (Francia et al., 2012). Alternatively, evidence exists that endogenous RNA transcripts may serve as templates for HR repair, and loss of RBPs may have a primary effect on these template RNA molecules (Keskin et al., 2014). Furthermore, if RBP-deficient HR defective tumors utilize Rad52 for repair as has been documented following loss of canonical HR repair genes such BRCA1, BRCA2, PALB2, and RAD51C (Rossi et al., 2021). Further mechanistic insight into how RBPs and other novel HRD drivers modulate HR repair will advance our understanding of the diverse mechanisms used to promote genomic stability in human cells.

While PARP inhibitors are gaining FDA approval in a variety of contexts. However, they are most commonly used for treating breast and gynecological cancers, and most on-going PARP inhibitor clinical trials are performed within this context. Across TCGA, we found that men tended to have higher HRD scores, strongly observed in the context of chromophobe kidney cancer, acute myeloid leukemia, pancreatic cancer, head and neck squamous cell carcinoma, and esophageal carcinoma. It is possible that because these cancers are enriched for HRD driven by previously undocumented RBPs that are likely to drive HRD, men with these specific HRD positive cancers may constitute an understudied population. Loss of RBPs may promote sensitivity to PARPi directly via induction of HRD, through secondary mechanisms such as induction of R-loops, or a combination thereof. Use of the HRD score and/or screening for mutations in the novel HRD driver genes we identified may provide biomarkers that could be used to stratify patients based on their predicted response to PARP inhibitors and expand the number of cancers that might benefit from such treatment. Alternatively, pharmacological induction of HRD by inhibiting RBPs may offer a novel approach to sensitize tumor cells to PARP inhibitors.

The novel drivers of HRD identified here were enriched for loci associated with cancer risk in prior GWAS studies. Risk-reduction surgery for BRCA-related breast cancer can decrease the risk of developing cancer from 85%-100%, but these surgeries are not without potential risk and should be focused on high-risk individuals (Nelson et al., 2014). Integration of GWAS studies with mechanistic understanding can increase the confidence in the relevance of GWAS-uncovered candidate cancer risk loci. In turn, this could advance cancer prevention programs by improving the genetic information available to genetic counselors and patients.

## MATERIALS AND METHODS

Further information and requests for resources and reagents should be directed to and will be fulfilled by the Lead Contact, Nidhi Sahni.

### Data sets utilized and data availability

TCGA RNAseq gene expression, alternative splicing, methylation, copy number, RPPA, and clinical data were downloaded from the GDC data commons (https://portal.gdc.cancer.gov/). TCGA CPTAC data were acquired from the manuscript’s supplemental information (Mertins et al., 2016). ICGC data were downloaded from the ICGC data portal (https://dcc.icgc.org/). GWAS data were downloaded from GWASdb v2 (http://jjwanglab.org/gwasdb) (Li et al., 2016). MicroRNA targets were downloaded from miRDB (http://mirdb.org/) (Wong and Wang, 2015). Table S1 contains all HRD tumors, with identified HRD drivers and gene annotations. RNAseq data generated in this manuscript are available at NCBI GEO accession GSE153396.

### Calculation of HRD score

We calculated the genomic scar HRD scores from SNP arrays following the previous published 3 components of HRD score: loss of heterozygosity (LOH) (Abkevich et al., 2012), large scale transitions (LST) (Popova et al., 2012), and telomere allelic imbalance (TAI or NtAI) (Birkbak et al., 2012), and used the sum of all 3 scores as the final HRD score (Knijnenburg et al., 2018; Marquard et al., 2015; Telli et al., 2016).

### Association of molecular data with HRD score

To analyze RNA, miRNA, and proteins that were associated with high HRD scores we performed regression with a generalized linear mixed effects model, taking the tumor type as a random effect. For gene expression changes from RNAseq data, the resulting regression coefficients were used for gene set enrichment analysis as described (Subramanian et al., 2005). For protein data from RPPA, regression coefficients and corresponding P values were reported directly with false discovery rate determined with Benjamini–Hochberg procedure. For miRNA data, regression coefficients and corresponding P values were reported directly with the FDR determined using Storey’s method. To define those pathways that were preferentially suppressed by miRNAs, we retrieved predicted miRNA gene targets from MicroRNA Target Prediction Database (miRDB). For each significant microRNA, the regression coefficient was added to all candidate target genes, yielding a positive score for genes predicted to be preferentially targeted by microRNAs in HRD tumors. Resulting microRNA scores were utilized for GSEA, and the top 5 significant pathways were reported.

### Threshold score to separate HRD positive vs. negative

To define a gold standard by which to define a threshold to identify HRD positive tumors, we selected breast, ovarian, prostate, and pancreatic tumors that had germline mutations in *BRCA1* or *BRCA2*. For determining the optimal threshold value, we performed stochastic sub-sampling using 50% of all patients for 1000 iterations. For each iteration, we determined a modified Youden’s J statistic defined as J’ = 2*true positive rate + true negative rate – 2, with increased weighting on the true positive rate because the definition of true positives was more robust than that of true negatives. The modal threshold value was determined to be 31, which was used to classify tumors as HRD positive or HRD negative.

### Identification of novel candidate HRD drivers

To identify candidate HRD drivers, we initially used canonical DDR genes (Lord and Ashworth, 2016) as well as highly validated candidates from two previous studies (Adamson et al., 2012; Herr et al., 2015). Genetic events were defined as either mutations or methylation events. Mutation events were constrained to variant allele frequencies (VAF) of at least 0.1. Methylation events were only considered if in the given tumor type methylation significantly correlated with gene expression (r≤-0.25), and the specific tumor exhibited both methylation and down-regulation (1 standard deviation) of the candidate gene. To assign genes to individual tumors, a scoring metric of 2*(HRD score, upper tertile) + (HRD score, lower 20^th^ percentile) – 0.5*(Coefficient of variation) was used to rank genes, with the highest scoring gene being assigned as the cause for a given tumor. To identify novel candidate genes for the next iteration, we first computed an interaction score using a merged protein interaction list from both the BioPlex affinity purification-mass spectrometry network (Huttlin et al., 2015) as well as a literature curated network (Menche et al., 2015). For all genes, we determined the number of interactions with genes identified to be candidate drivers of HR defects, and then z-transformed these values. This z-normalized network score was averaged with a z-normalized score for induction of HR defects from two siRNA screens (Adamson et al., 2012; Herr et al., 2015). All genes that score above the 25^th^ percentile of candidate genes identified in the prior iteration were added for the next iteration. This process was completed until convergence, that is no additional candidate genes were identified.

### DR-GFP HR reporter assay

The DR-GFP reporter assay was performed in U2OS cells per previous publications (Pierce et al., 1999). Transfection with siRNA or the RFP-tagged protein of interest was performed 12 hours prior to transfection with I-SceI (Addgene #26477) or GFP control (Addgene #89684). All transfections were performed with Lipofectamine 3000 per the manufacturer’s instructions. Two days after I-SceI/GFP transfection, samples were analyzed by flow cytometry. After gating for singlets and cell size, GFP positivity was gated based on SSC vs. GFP intensity. All values were normalized to the average of siCTRL and non-transfected controls for siRNA experiments, and RFP-tagged wild-type protein overexpression for analysis of RFP-tagged mutant proteins.

### RFP-tagged mutant constructs

To generate point mutations, we implemented a modified high-throughput site-directed mutagenesis pipeline described previously (Sahni et al., 2015; Yi et al., 2017). Briefly, we used the corresponding wild-type reference ORFs from their Entry clones in human ORFeome as template for a 3-step PCR experiment. For a given mutation, PCR cloning consisted of two “primary PCRs” to generate gene fragments, and one “fusion PCR” to obtain the mutated ORF. For the primary PCRs, two universal primers, Tag1-M13F and Tag2-M13R (sequences shown below), and two ORF-specific internal forward and reverse primers were employed. The two universal primers allowed the preservation of the attL sites on both ends of the ORF. The mutation-specific primers (namely MutF and MutR), encompassing the desired single nucleotide change, were designed to have an overlapping region of ∼40 base pairs. The two ORF fragments flanking the mutation of a gene were amplified using the primer pair Tag1-M13F and MutR, and the primer pair Tag2-M13R and MutF, respectively. For the fusion PCR, the two primary PCR fragments were fused together using the primer pair Tag1 and Tag2 (sequences shown below) to generate the single amino acid change mutation allele. The final product was a full-length ORF harboring the desired mutation. All wild-type and mutant allele clones were transferred by Gateway recombination into a mammalian expression vector containing a C-terminal RFP tag. For subsequent sequence confirmation, the inserts were PCR amplified with KOD HotStart Polymerase (Novagen) and verified by Sanger sequencing.

Primer sequences:

- *Tag1-M13F:* 5’-*GGCAGACGTGCCTCACTCCCAGTCACGACGTTGTAAAACG-3’*
- Tag2-M13R: 5’-CTGAGCTTGACGCATTGCTAGTGTCTCAAAATCTCTGATGTTAC-3’
- Tag1: 5’-GGCAGACGTGCCTCACTACT-3’
- Tag2: 5’-CTGAGCTTGACGCATTGCTA-3’

### IR induced foci formation

Cells were grown on glass coverslips, irradiated with either 5 Gy or 10 Gy, and incubated as specified. To analyze chromatin bound fractions, the soluble fractionwas extracted prior to fixation, as described (McGrail et al., 2017). For analysis of RFP-tagged protein foci formation, cells were transfected 48 hours prior to irradiation using Lipofectamine 3000 per manufacturer’s instructions prior to fixation and detection of phosphorylated histone variant H2AX using indirect immunofluorescence with anti-γH2AX (clone JBW301, Millipore Sigma). For IR-induced Rad51 foci, cycling cells were pulse labeled with 10 μM EdU prior to irradiation. EdU was labeled with CLICK chemistry as described (Fang et al., 2019). Nuclei and Rad51 foci were segmented, and EdU positivity was determined from integrated intensity in each nuclei compared to a no EdU stained control. Rad51 positivity was analyzed only in EdU positive cells, and defined relative to control cells without irradiation. For siRNA experiments, cells were transfected with siRNA 48 hours prior to irradiation. For Pladienolide B experiments, cells were treated with Pladienolide B (Tocris) 24 hours prior to irradiation. Cells were imaged by fluorescence microscopy (Eclipse TE2000E, Nikon), capturing all images for a given replicate simultaneously to assure no variances in light intensity. All quantification was performed in Matlab R2016a.

### PARP sensitivity assays

For short term assays, cells were plated in 96 well plates before treatment with specified concentrations of BMN673 (Selleck), AZD2281 (Selleck), Pladienolide B, or vehicle control. Cells were incubated for 5 days, and viability was assessed using PrestoBlue (Invitrogen) relative to vehicle controls (DMSO for BMN673 and AZD2281, ethanol for Pladienolide B). Synergy was assessed using the Chou-Talalay combination index (Chou, 2006). For clonogenic assays, cells were plated in 12 well plates and incubated with drugs for two weeks before fixation and staining with crystal violet. Plates were scanned, and the crystal violet was extracted with 10% acetic acid. Absorbance of solubilized crystal violet was measured at 590 nm using a plate reader, and viability was normalized to a solvent-treated control. For stable shRNA cells, Dharmacon pGIPZ Lenti shRNA vectors were used including a pGIPZ non-silencing control (RHS4346), shDDX3X (V3LHS_644473, V2LHS_202531), shSF3B3 (V3LHS_644840, V2LHS_43924), and shBRCA2 (V2LHS_89238, V2LHS_89237).

### RNAseq with depletion of RNA binding proteins

Cells (BT-549, MDA-MB-231, or U2OS) were transfected with the desired siRNAs (Table S3) using Lipofectamine 3000 48 hours prior to RNA isolation with a Qiagen RNeasy Kit. RNA quality was confirmed using the Agilent TapeStation RNA reagents according to the manufacturer’s protocol. Libraries were prepped using the Lexogen QuantSeq 3′ mRNA-Seq Library Prep Kit FWD for Illumina with 6nt unique dual indexing using. 100ng of RNA was used as input material for an automated protocol adapted for the Perkin Elmer SciClone NGS Workstation. Libraries were analyzed for quality using the Agilent TapeStation High Sensitivity DNA kit and for quantity using Qubit dsDNA HS assay, in 384-well format, using 20µL reactions in triplicate (19µL working reagent + 1µL sample or standard) with an 11-point standard curve from 0-10ng/µL. Plates were shaken in the plate reader for 5 seconds, then measured for fluorescence (excitation: 480nm, emission: 530nm). Sample concentrations were determined using the standard curve. Libraries made from each RNA sample were then pooled at 25 nM each, denatured with 1M NaOH added to a 0.2M final concentration (5 min at room temperature), and quenched with 200mM Tris HCl (pH 7). 1% PhyX spike-in (Illumina) was added then pooled, denatured libraries were run on an Illumina NovaSeq with a NovaSeq 6000 SP Reagent Kit (100 cycles) using 51bp reads, 6bp index reads, and paired-end single read parameters.

### Identification of RNA binding proteins that may modulate DDR genes

To test whether RBPs were modulating DDR genes, we performed multiple analyses. First, RNAseq analysis following siRNA-mediated depletion of 17 RBPs in 3 cell lines was used to identify differential DDR gene expression relative to either siCTRL, or siBRCA1/siBRCA2 by paired t-test. Next, TCGA tumors were analyzed to detect decreased DDR gene expression relative to HR competent tumors or tumors with HRD caused by DDR genes using a generalized linear mixed effects model, taking the tumor type as a random effect. Finally, the same comparisons in TCGA tumors was made using alternative splicing instead of gene expression levels. We considered gene expression changes that caused decreased gene expression with an FDR of less than 10%, or events that were detected in both patient tumors and cell lines with a nominal P value of at least 0.05. For alternative splicing, increased or decreased alternative splicing at 10% FDR was considered an event. The resulting network was visualized in Cytoscape v3.5.1.

### Statistics

Specific statistical tests are discussed within corresponding sections. In general, pan-cancer associations with HRD score were determined using a generalized linear mixed effects model, taking the tumor type as a random effect. Multiple comparisons were corrected for using either Storey’s method (large number of variables) or the Benjamini–Hochberg procedure (smaller number of variables). Comparisons of normally distributed data were made using either t tests (2 groups) or one-way ANOVA (3+ groups) with appropriate post-hoc analysis. Comparisons of non-normally distributed data were made using rank sum test (2 groups) or Kruskal-Wallis (3+ groups) with appropriate post-hoc test. Correlations were assessed using Spearman rank correlation coefficients. Categorical variables were compared using Fisher’s exact test.

## Supplemental Tables

Table S1. Identified causes of HR defects, Related to Figure 3.

Table S2. Functional analysis of putative novel drivers of HRD, Related to Figure 4.

Table S3. siRNAs used in this study, Related to Materials and Methods

## ACKNOWLEDGEMENTS

N.S. is a CPRIT Scholar in Cancer Research with funding from the Cancer Prevention and Research Institute of Texas (CPRIT) New Investigator Grant RR160021. NS is a recipient of the Liz Tilberis Early Career Award funded by Ovarian Cancer Research Alliance grant #649968 and Young Investigator grant from the Breast Cancer Alliance. D.J.M. was supported by Susan G. Komen PDF17483544 and NCI grant K99CA240689. S.Y. was supported by Komen Foundation grant CCR19609287. M.L.M was supported by the Susan G. Komen Foundation (CCR17488145), National Cancer Institute of the NIH (R00CA175293), the Kimmel Scholar (SKF-16-135) and Lynn Sage Scholar awards. Additional support was provided by a Department of Defense Era of Hope Scholar Award (W81XWH-10-1-0558) and George and Barbara Bush Endowment for Innovative Cancer Research to S.-Y.L. and U01CA217842 to G.B.M. We appreciate MD Anderson Cancer Center core facilities funded by grant CA016672: the Functional Genomics Core (shRNA and ORFeome Core) for reagents and technical assistance and the Characterized Cell Line Core for STR DNA fingerprinting and mycoplasma testing. The results here are in whole or part based upon data generated by the TCGA Research Network: http://cancergenome.nih.gov/.

## Author Contributions

D.J.M., S.Y., and N.S. conceived the study; D.J.M wrote the manuscript with significant input from B.D., M.L.M, G.B.M., S.Y. and N.S. D.J.M and B.F. performed computational analysis with input from G.B.M., S.Y. and N.S. D.J.M. conducted most of the experiments with help from N.S., R.S.S., Yang Li and L.H. N.S., Yongsheng Li., and L.H. generated the allele libraries. N.S., R.S.S., and Yang Li performed siRNA treatment, library preparation and RNA sequencing. S.-Y.L., S.Y., and N.S. provided intellectual input and supervision throughout the course of the study. All authors read and approved the final manuscript.

## SUPPLEMENTAL FIGURES

**Figure S1.**
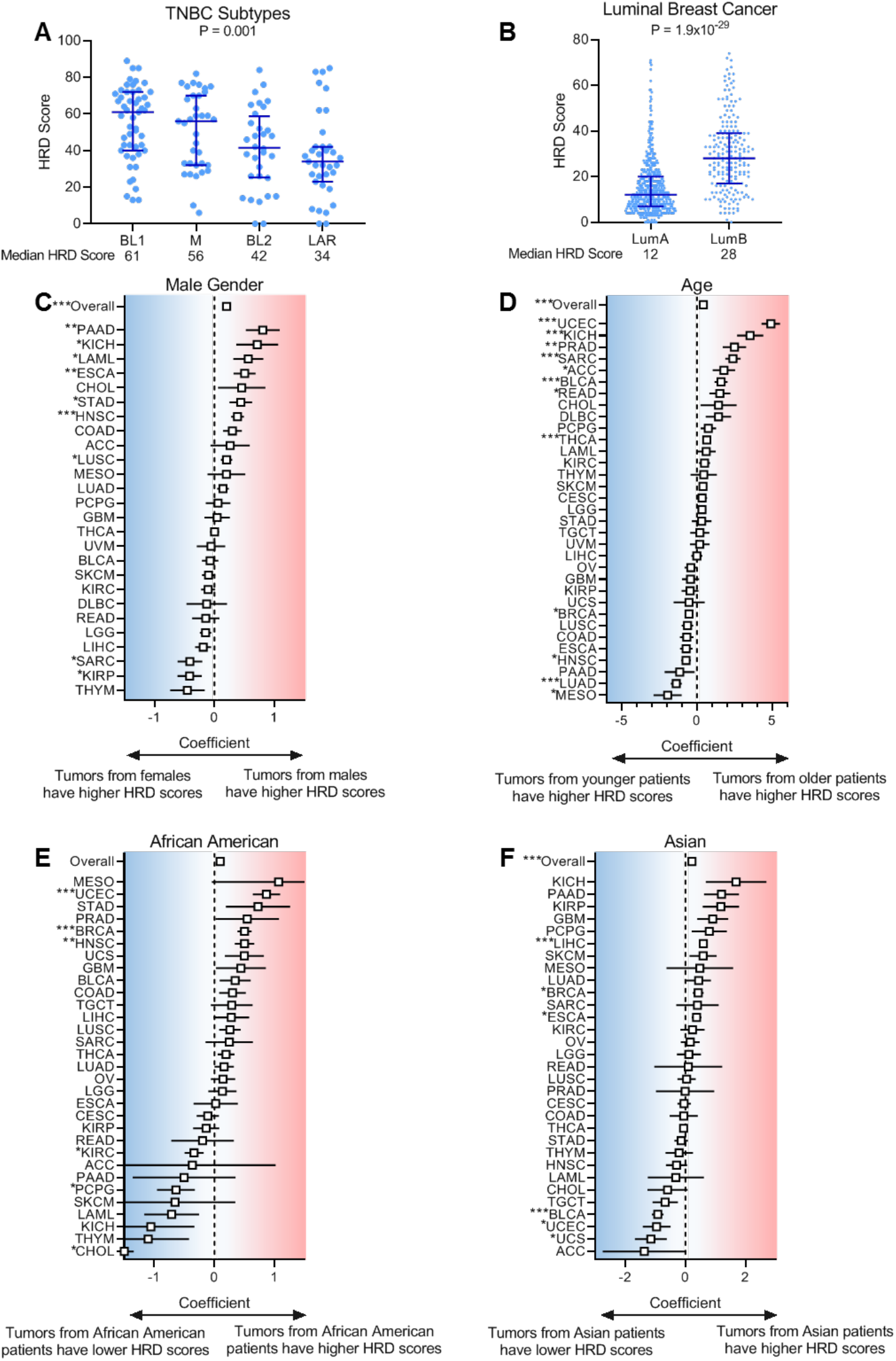
Demographics of HR defects, Related to Figure 1. **(A)** HRD score for specific TNBC subtypes, shown as median with interquartile range. Basal-like 1 (BL1), metastatic (M), basal-like 2 (BL2), luminal-androgen receptor (LAR). Kruskal-Wallis test. **(B)** HRD score in luminal A (LumA) and luminal B (LumB) breast cancer, shown as the median with interquartile range. Rank-sum test. **(C-F)** Regression coefficients for HRD score with the indicated demographic information. The overall regression coefficient was determined using a generalized, linear mixed model with tumor type as a random effect. (C) Positive coefficient indicates HRD score is higher in tumors from male patients with cancer. (D) Positive coefficient indicates HRD score increases in tumors from patients of older age. (E) Positive coefficient indicates HRD score is increased in tumors from African American patients with cancer. (F) Positive coefficient indicates HRD score is increased in tumors from Asian patients with cancers. *p<0.05, **p<0.01, ***p<0.001.

**Figure S2.**
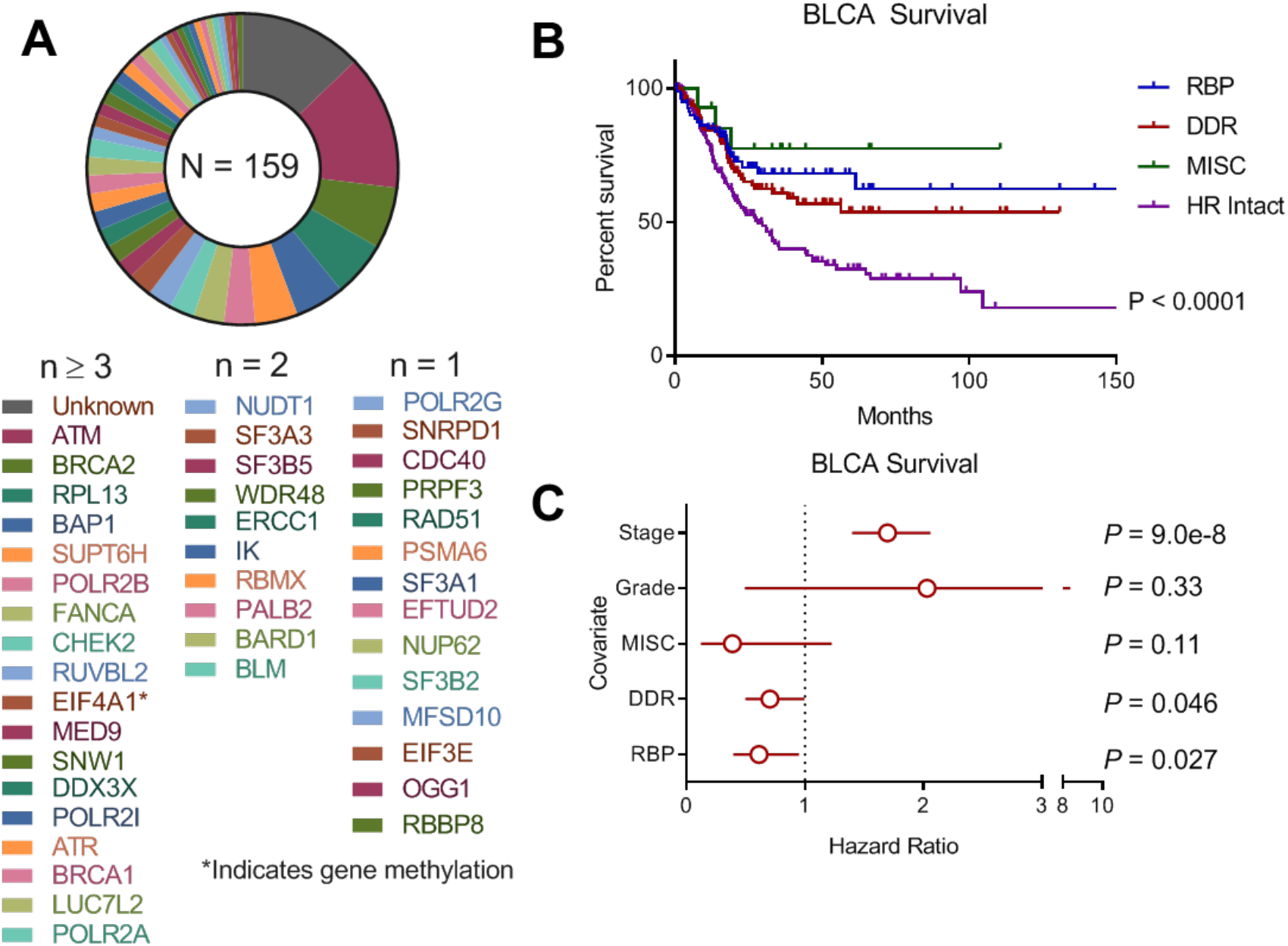
Survival of patients with bladder cancer by type of HR defect, Related to Figure 3. **(A)** Genes identified as putative causes of HR defects in tumors from patients with bladder cancer. **(B)** Kaplan-Meier curves showing overall survival of bladder cancer patients, stratified by general type of putative HR defect compared to patients with HR competent tumors. Log-rank test to assess survival differences. **(C)** Multivariate survival analysis with Cox proportional hazards model.

**Figure S3.**
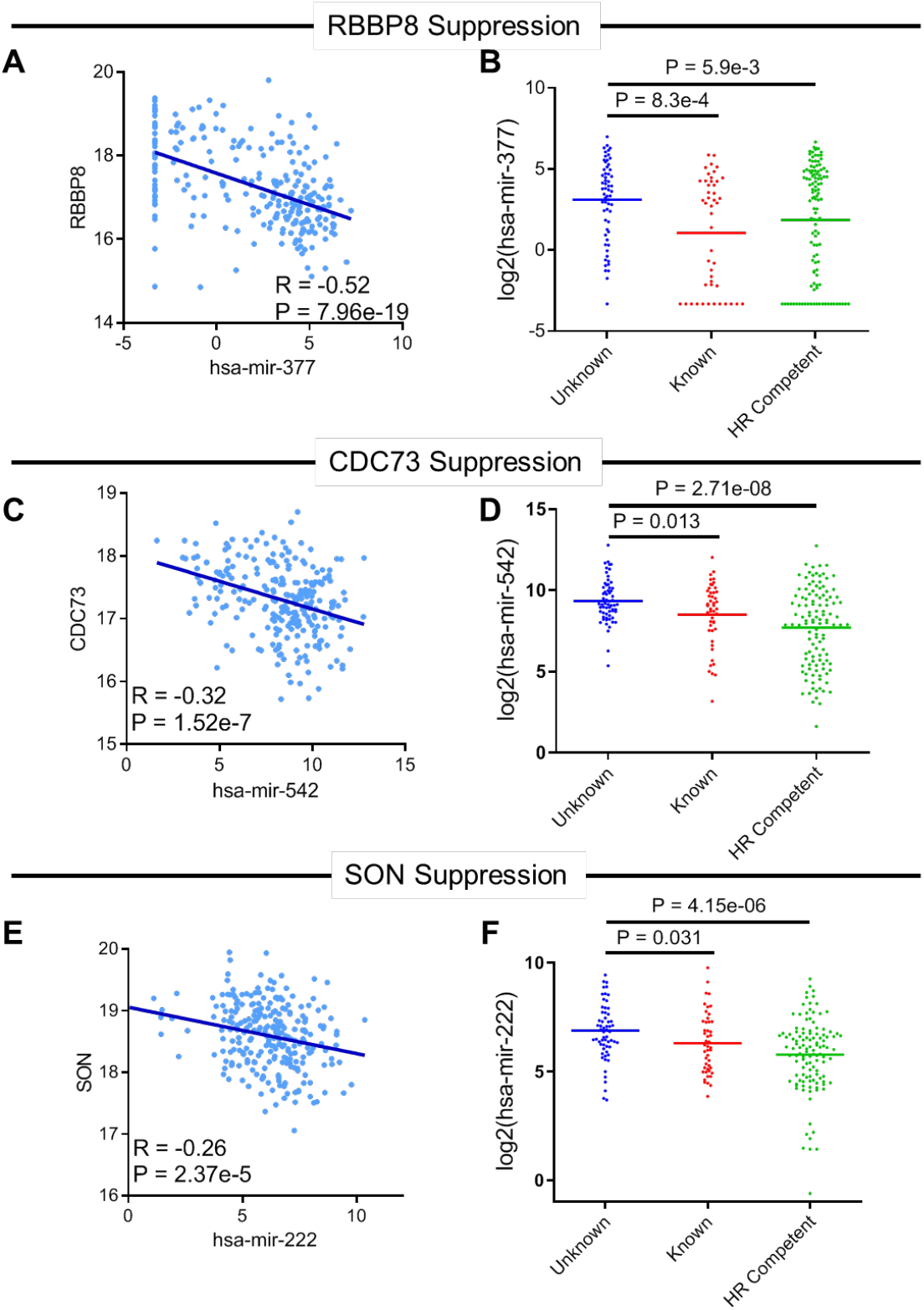
MicroRNA control of HR defects in sarcoma, Related to Figure 3. **(A)** Scatter plot showing an inverse correlation between RBBP8 and hsa-mir-377 expression. Spearman correlation coefficient. **(B)** Differential expression of hsa-mir-377 in tumors with HRD of unknown cause, HRD in which a candidate driver could be identified, and HR-competent tumors. Kruskal-Wallis with Dunn’s posthoc test. **(C)** Scatter plot showing an inverse correlation between CDC73 and hsa-mir-543 expression. Spearman correlation coefficient. **(D)** Differential expression of hsa-mir-543 in tumors with HRD of unknown cause, HRD in which a candidate driver could be identified, and HR-competent tumors. Kruskal-Wallis with Dunn’s posthoc test. **(E)** Scatter plot showing an inverse correlation between SON and hsa-mir-222 expression. Spearman correlation coefficient. **(F)** Differential expression of hsa-mir-222 in tumors with HRD of unknown cause, HRD where a candidate driver could be identified, and HR-competent tumors. Kruskal-Wallis with Dunn’s posthoc test.

**Figure S4.**
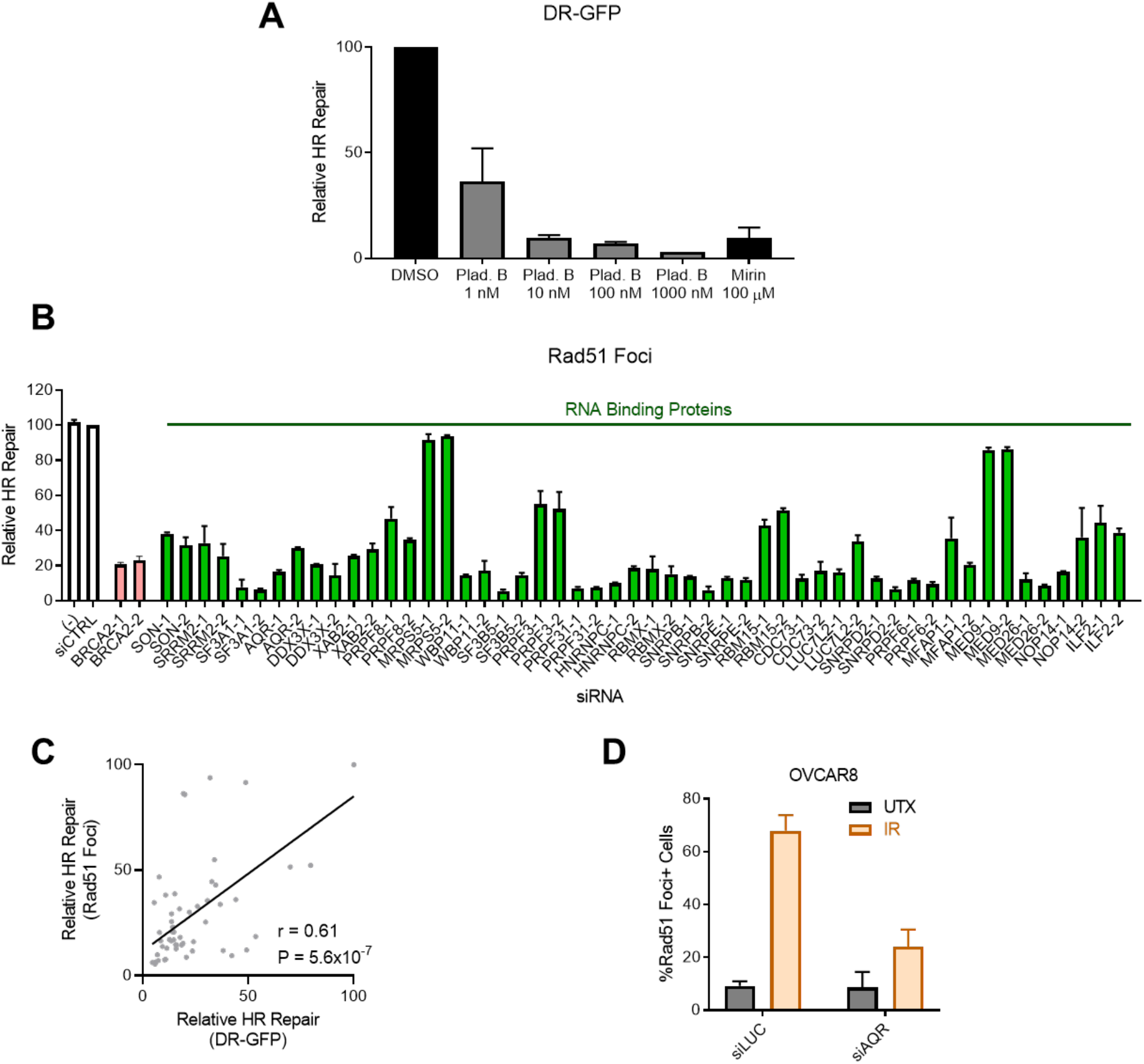
HR defects associated with loss of RBPs, Related to Figure 3. **(A)** A DR-GFP HR reporter system was used to measure HR in U2OS cells treated with the indicated concentrations of the spliceosome inhibitor pladienolide B, the Mre11–Rad50–Nbs1 (MRN) inhibitor Mirin, or a DMSO vehicle control. **(B)** Quantification of IR-induced Rad51 foci in cycling (EdU^+^) U2OS cells following siRNA-mediated depletion of the indicated RBPs. Reported as repair relative to control siRNA (siCTRL). N = 2 per cell line. **(C)** Correlation between relative HR repair by DR-GFP (Figure 4A) and by Rad51 foci in EdU^+^ cells (Figure S4B). Pearson correlation coefficient. **(D)** Quantification of IR-induced Rad51 foci in cycling (EdU^+^) OVCAR8 ovarian cancer cells following siRNA-mediated depletion of AQR or siLuciferase (siLUC) control. Reported as repair relative to control siRNA. N = 2 per cell line.

**Figure S5.**
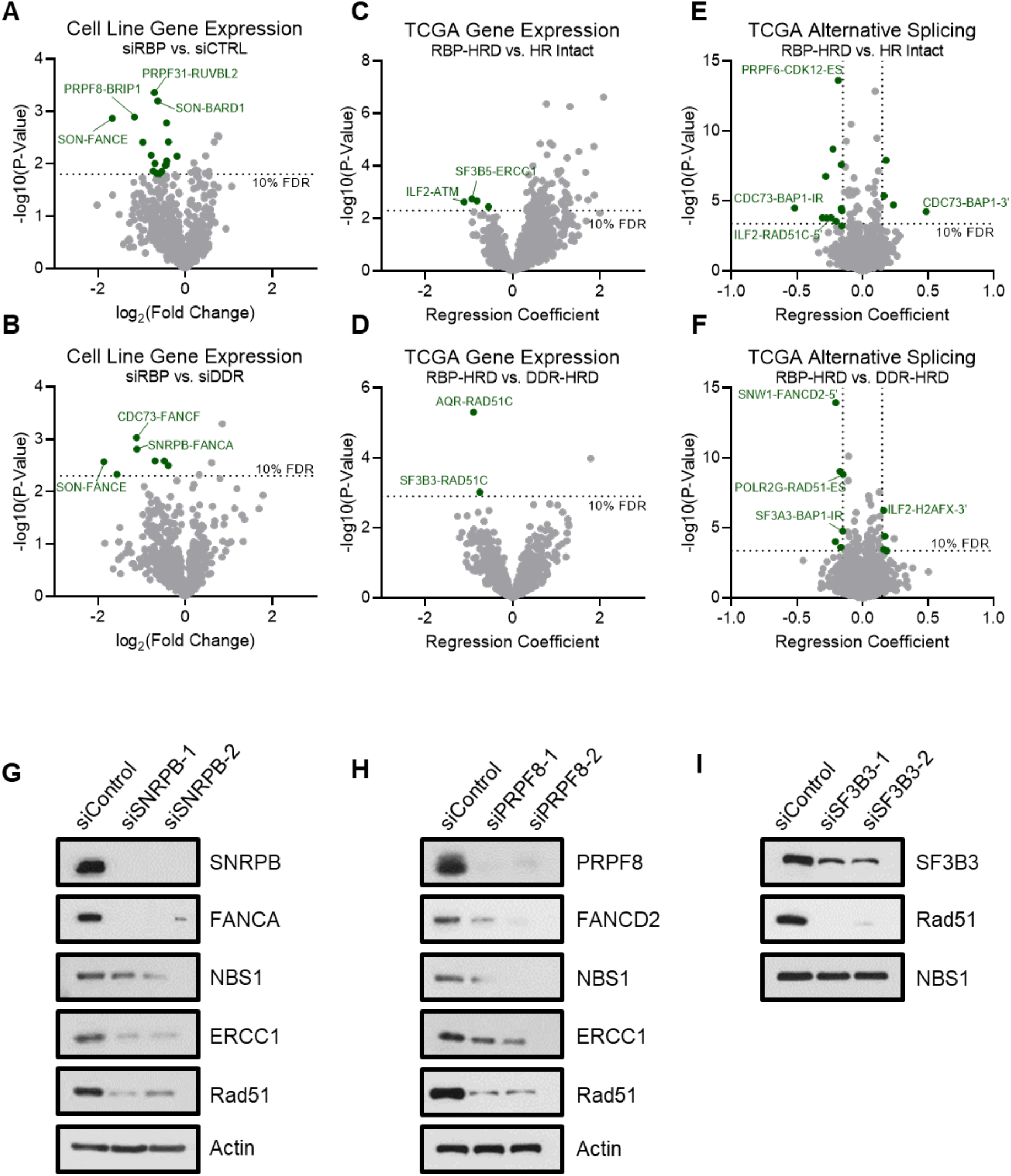
Control of DDR gene expression and splicing by RBPs, Related to Figure 5. Volcano plots showing genes that are either suppressed or alternatively spliced upon RBP loss. Genes reaching statistically significant changes are highlighted in green. Annotations are listed as “RBP Gene”-“DDR Gene,” for example, SON-FANCE in panel A indicates SON loss reduces expression of FANCE. **(A)** DDR gene expression in cells with RBP expression knocked down using siRNA (siRBPs) compared with an siRNA negative control (siCTRL) in U2OS, MDA-MB-231, and BT-549 cell lines. RNA-seq was performed in duplicate for each cell line and averaged after log transformation. Significance was determined using a paired t-test with the FDR determined by the method of Storey. **(B)** DDR gene expression in cells with RBP expression knocked down using siRNAs (siRBPs) compared with knock-down of BRCA1 and BRCA2 with siRNA (siDDR) in U2OS, MDA-MB-231, and BT-549 cell lines. RNA-seq was performed in duplicate for each cell line and averaged after log transformation. Significance was determined using a paired t-test with the FDR determined by the method of Storey. **(C)** DDR gene expression in TCGA patient tumors with HRD attributed to loss of RBPs compared to HR competent-patient tumors using a generalized linear mixed model with tumor type as a random effect. The FDR was determined by the method of Storey. **(D)** DDR gene expression in TCGA patient tumors with HRD caused by RBP aberrations (mutation or methylation) compared to HRD caused by aberrations DDR gene aberrations determined using a generalized linear mixed model with tumor type as a random effect. The FDR was determined by the method of Storey. **(E)** Alternative splicing of DDR genes in TCGA patient tumors with HRD putatively caused by RBP aberrations (mutation or methylation) compared to tumors from HR-competent patients using a generalized linear mixed model with tumor type as a random effect. The FDR was determined by the method of Storey. **(F)** Alternative splicing of DDR genes in TCGA patient tumors with HRD putatively caused by RBP aberrations (mutation or methylation) compared to HRD caused by DDR gene aberrations determined using a generalized linear mixed model with tumor type as a random effect. The FDR was determined by method of Storey. **(G)** Validation of depletion of SNRPB causing loss of canonical HR genes FANCA, NBS1, ERCC1, and Rad51 identified in Figure 6A/S5A-F at the protein level by western blot in BT-549 triple-negative breast cancer cells. **(H)** Validation of depletion of PRPF8 causing loss of canonical HR genes FANCD2, NBS1, ERCC1, and Rad51 identified in Figure 6A/S5A-F at the protein level by western blot in BT-549 triple-negative breast cancer cells. **(I)** Validation of depletion of SF3B3 causing loss of canonical HR gene Rad51 identified in Figure 6A/S5A-F at the protein level by western blot in BT-549 triple-negative breast cancer cells.

**Figure S6.**
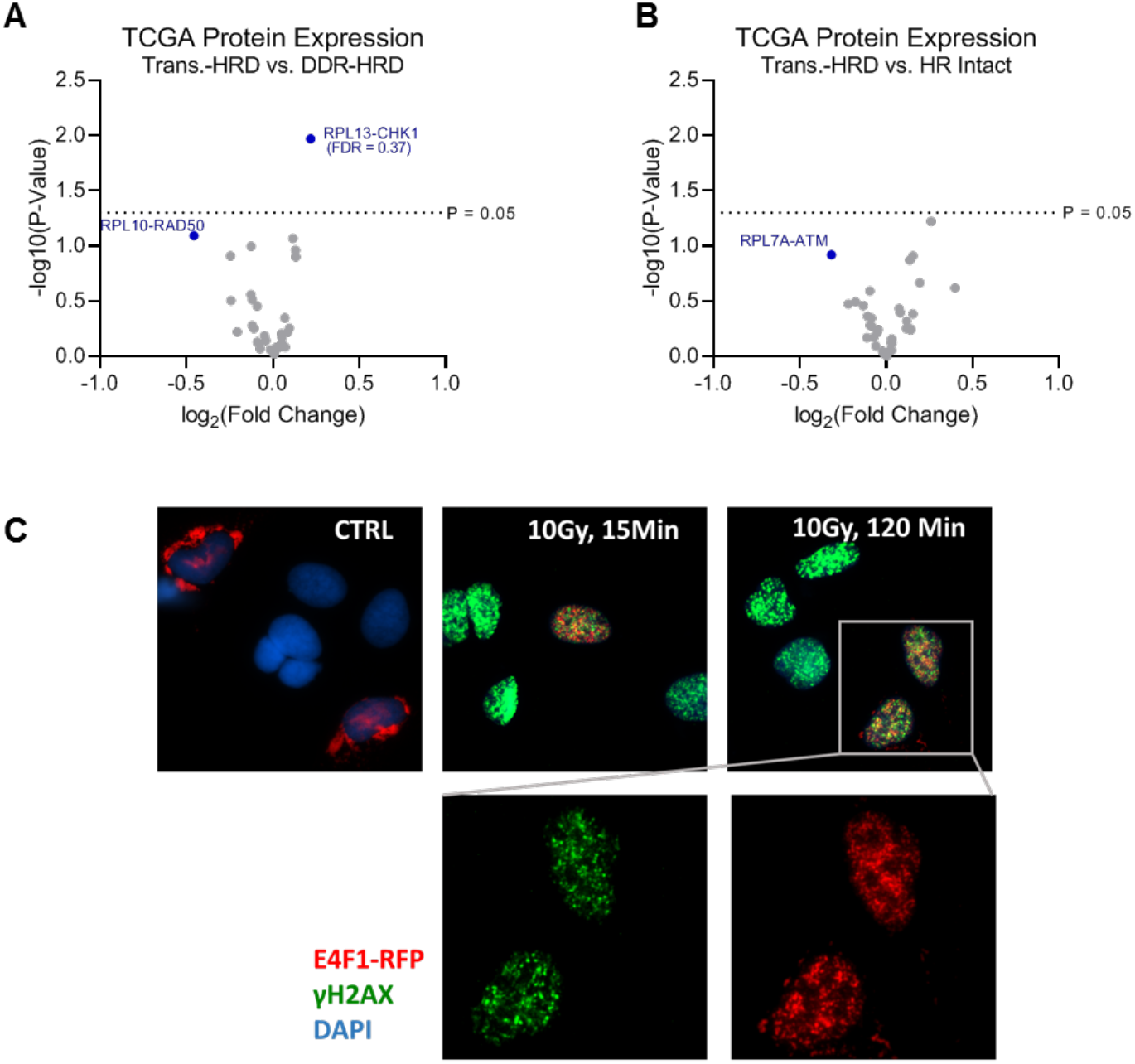
Control of DDR protein expression by translation genes, Related to Figure 5. **(A)** DDR protein expression based on RPPA data for TCGA patient tumors with HRD putatively caused by aberrations in translation genes (Trans.-HRD) compared to tumors from HR-competent patients using a generalized linear mixed model with tumor type as a random effect. FDR determined by Benjamini–Hochberg procedure. Annotations are listed as “Translation Gene”-“DDR Gene”. **(B)** Comparison of DDR protein expression based on RPPA data for TCGA patient tumors with HRD putatively caused by aberrations (mutation or methylation) in translation genes compared to HRD caused by aberrations in DDR genes determined using a generalized linear mixed model with tumor type as a random effect. FDR determined by Benjamini–Hochberg procedure. Annotations are listed as “Translation Gene”-“DDR Gene”. **(C)** IR-induced foci of translation factor E4F1-RFP, with co-staining for DNA double-strand break marker γH2AX. Soluble protein was extracted prior to fixation.

